# High-throughput Single-Virion DNA-PAINT Reveals Structural Diversity, Cooperativity and Flexibility during Selective Packaging in Influenza

**DOI:** 10.1101/2025.02.11.637713

**Authors:** Christof Hepp, Qing Zhao, Nicole Robb, Ervin Fodor, Achillefs N. Kapanidis

## Abstract

Influenza A, a negative-sense RNA virus, has a genome that consists of eight single-stranded RNA segments. Influenza co-infections can result in re-assortant viruses that contain gene segments from multiple strains, causing pandemic outbreaks with severe consequences for human health. The outcome of re-assortment is likely influenced by a selective sequence-specific genome packaging mechanism.

To uncover the contributions of individual segment pairings to selective packaging, we set out to statistically analyse packaging defects and inter-segment distances in individual A/Puerto Rico/8/34 (H1N1) (PR8) virus particles. To enable such analysis, we developed a multiplexed DNA-PAINT approach capable of assessing the segment stoichiometry of more than 10,000 individual virus particles in one experiment; our approach can also spatially resolve the individual segments inside complete virus particles with a localisation precision of ~10 nm.

Our results show the influenza genome can be assembled through multiple pathways in a redundant and cooperative process guided by preferentially interacting segment pairs and aided by synergistic effects that enhance genome assembly, driving it to completion. Our structural evidence indicates that the interaction strength of segment pairs affects the spatial configuration of the gene segments, which appears to be preserved in mature virions. As our method quantified the interactions of whole influenza segments instead of identifying individual sequence-based interactions, our results can serve as a template to quantify the contributions of individual sequence motifs to selective packaging.

## INTRODUCTION

Influenza A is a negative-sense, single-stranded RNA virus with a genome that consists of eight RNA segments. Each segment forms a ribonucleoprotein complex (vRNP) with many copies of virus-encoded nucleoprotein (NP) and three subunits of the viral RNA polymerase (PB1, PB2 and PA) [1]. Each vRNP is organized into a highly flexible double-helical structure made up from NP, with the polymerase situated at one end [2, 3]. The RNA binds to a groove on NP that is solvent-exposed in the mature vRNP structure [4–6].

Influenza causes pandemics with severe, worldwide consequences for human health, such as the Spanish Flu of 1918. Pandemics occur whenever zoonotic influenza viruses gain the ability for human-to-human transmission through genetic variation. In general, co-infection of the same host with multiple influenza strains can lead to novel virus strains; such strains are generated by genetic re-assortment, which in turn relies on co-packaging of gene segments from different strains [7]. Hence, an important strategy for controlling future influenza epidemics and pandemics is to gain the ability to predict the probability of genetic reassortments; such a prediction requires detailed understanding of the molecular mechanisms of the viral segment packaging during the assembly of a virus particle.

However, despite the importance of vRNP packaging for the life cycle of influenza, the packaging mechanism remains puzzling. After influenza vRNPs have been synthesized in the nucleus, they are transported to the cytoplasmic membrane via Rab11 and attach to the inside of budding virus particles, likely binding M1 [8, 9]. The segmented viral genome is thought to be assembled inside the cytoplasm. Recent evidence favours a selective model over stochastic assembly, postulating the presence of packaging signals that control the incorporation of a complete set of influenza segments into virions [10, 11]. A few key terminal sequence motifs that support the incorporation of influenza segments into virus particles have been identified through bioinformatic approaches and mutational analysis [12–21]. In some instances, packaging defects in one segment affected incorporation rates for the other segments [22–26]. Sequence motifs that interact with each other in selective packaging have also been identified [27–29].

In contrast, other studies, based mainly on biochemical approaches [sequencing of psoralen crosslinked, ligated and selected hybrids (SPLASH), dual crosslinking, immunoprecipitation, and proximity ligation (2CIMPL) and psoralen analysis of RNA interactions and structures (PARIS)] predict a complex and extensive network of intersegment contacts along the entire length of the gene segments [30–34]. However, recent data questioned the role of SPLASH-identified RNA-RNA contacts in the packaging process, illustrating the need for their cross-validation by independent methods [35].

An alternative strategy for identifying packaging motifs and pathways is to quantify the interactions between individual segment pairs by structurally analysing the influenza genome in individual virus particles. Electron tomographs of individual virions showed that the packaged vRNPs align in a parallel fashion during virus budding, with a centrally placed vRNP contacting all other vRNPs surrounding it (adopting a “7+1” configuration), with recent EM data revealing that at least a part of the mature virus population keeps the same arrangement [36–40]. A tentative, partial identification of the segments based on vRNP length pointed towards a varying segment arrangement within the 7+1 configuration [38]. Further, some EM data suggest that there are regions where individual vRNPs contact each other [38]. However, the EM-based approach for evaluating sequence-based information on the packaging process (see above) is limited by two major drawbacks: 1) the identification of all eight segments inside a single virus particle is technically challenging and has not been realized so far, and 2) the throughput of electron tomography is too low to draw definitive conclusions from the observed configurational complexity.

The co-localization of gene segments inside virus particles or as a part of intracellular, ternary segment complexes can also provide clues on segment interactions during the packaging process. Several studies on selective packaging in influenza virus make use of the detection of single influenza gene segments by fluorescence-in-situ hybridization (FISH), analogous to single-molecule RNA (smRNA) FISH in eukaryotic cells [11, 41–45]. Based on the co-localization frequency of selected gene segments in infected cells, likely sequences of vRNP association events that lead to the formation of specific subcomplexes during cytoplasmic transport were predicted [46]. Recently, an smRNA FISH approach was able to sequentially interrogate all eight gene segments in infected cells, thus directly visualizing a plethora of multi-segment subcomplexes, some of which were twice as frequent as any other subcomplex of the same rank [47]. However, the involved sequential applying and stripping of individually prepared FISH probe sets is time-consuming and may denature and perturb the sample too much to permit structural studies. Moreover, most experiments focused on vRNPs inside infected cells, where they may co-localize for reasons other than being part of the same subcomplex, e.g., during vesicular co-transport [42, 46, 47].

Consequently, describing the interactions between the segments in a reliable and quantitative fashion requires a methodology that can rapidly examine a large number (tens of thousands) of individual mature virus particles, can detect *every* gene segment within them, and (ideally) permit spatial analysis of the segment configuration. A powerful method that can fulfil the above requirements is DNA-PAINT (DNA-based points accumulation for imaging in nanoscale topography), a super-resolution microscopy method for optically reconstructing biomolecular assemblies from the localization of transient hybridization events to oligonucleotide sequences on a target structure [48, 49]. Using multiple target sequences (“barcodes”) permits the interrogation of various molecular targets at once by sequentially adding fluorescently labelled complementary oligonucleotides (multiplexing). Recently, multiplexed DNA-PAINT has been combined with *in situ* hybridization to elucidate the spatial organization of both DNA and RNA in individual eukaryotic cells [50–53].

To bridge the knowledge gap between genetically identifying packaging signals and testing their contribution through structural analysis, we developed a versatile multiplexed DNA-PAINT approach that detects the presence (or absence) of all eight viral segments inside of >10,000 individual virus particles per experiment, while spatially resolving individual segments inside complete virus particles with a precision of better than 10 nm. Our results show that the influenza genome assembly is a cooperative process, with a tendency towards virions with higher segment counts. Certain segment pairs co-appear preferentially, indicating the presence of segment-specific interactions; we also observed that all segments interacted to some extent. Both the measured inter-segment distances and the spatial distribution of segments inside virions suggest a flexible, but non-random genome arrangement that correlated with the detected segment pair frequencies. Our comprehensive description of the influenza genome structure can serve as the foundation for verifying the importance of packaging signal candidates identified by other complementary methods.

## METHODS

### Virus strain

A/Puerto Rico/8/34 (H1N1) influenza virus (PR8) purified from embryonated chicken eggs was purchased from Charles River Laboratories, aliquoted, flash-frozen in liquid N_2_ and stored at −80 °C before use.

### Probe design

Publicly accessible gene sequences for PR8 were used for the probe design (Pubmed EF467817.1 - EF467824.1, AF389122.1). Target sequences for barcoded hybridization probes on all eight gene segments were selected using Stellaris Probe Designer 4.2 from LGC Biosearch Technologies (Organism: Human, Masking Level: 5, Max. number of probes: 48, oligo length: 20, min. spacing length: 2). The sequences for the barcode extensions were derived from previously published DNA origami structures and consist of 5’-TT-3’ spacer at the 3’ end, followed by a 9-nt docking sequence [41]. All barcoded probes are summarized in Table S2. Fluorescent imager strands have been previously described, are summarized in Table S1 and were ordered from metabion international AG (Planegg, Germany) [41]. The fluorescent modification constituted a Cy3B modification at the 3’ end of every imager strand. Imager P6 was extended by 1 nt into the spacer (Table S1, underlined) to adjust the binding and dissociation kinetics to other imagers.

### Virus immobilization, preparation and hybridization

Glass Coverslips (Epredia, 24 x 60 mm, # 1.5) were sonicated in 2% Hellmanex III (Hellma Analytics, Germany) for 15 min to remove contaminations (45 kHz, 100 W). To remove Hellmanex, the coverslips were washed five times with MilliQ water, sonicated for 5 min and washed with MilliQ water another five times. The cleaned coverslips were dried under a nitrogen flow, plasma cleaned for 3 min at 100 W (Henniker Plasma, HPT-100) and immediately used for immobilisation.

Virus particles were thawed on ice. Depending on the application, particles were fixed before immobilisation by adding 16% formaldehyde (Thermo Scientific) to a final concentration of either 4% formaldehyde (high-throughput stoichiometry analysis) or 0.5% formaldehyde (super-resolution experiments) in HEPES-buffered saline (HBS), followed by an incubation at room temperature for 10 min and storage on ice until use. The virus aliquot was diluted 1:1000 in 0.9% NaCl and centrifuged for 2 min at 12,470 g to remove large aggregates. 10 µL of the dilution were added to a CultureWell gasket (Ø 6 mm, Grace Biolabs, USA) and evaporated at 40°C. After the sample had dried, the gasket was removed from the coverslip, a sticky slide VI^0.4^ (Ibidi, Germany) was mounted and the sample was washed with HBS.

Hybridization of the primary, barcoded probes was performed as previously described, with modifications [11, 54]. After fixing and immobilizing the sample as described above, the virus particles were permeabilized with 0.5% Triton X-100 in HBS for 15 min. After washing with 2x sodium citrate buffer (SSC), the sample was incubated with hybridization buffer HB (2x SSC, 10% formamide, 200 mg/mL E.coli tRNA, 20 mg/mL BSA, 1% RNasin Plus (Promega)) in the absence of probes for 10 min to block unspecific probe attachment. After blocking, equal volumes of the eight probe sets binding to the respective gene segments were added to a final total probe concentration 4 µM in buffer HB and incubated overnight at 37°C. After the hybridization, the unbound probes were removed by incubation with clearing buffer CB (2x SSC, 10% formamide) for at least 30 min until DNA-PAINT microscopy was performed.

### DNA-PAINT

All experiments were performed on a commercial fluorescence microscope (Nanoimager, Oxford Nanoimaging, UK). The sample was imaged using total internal reflection fluorescence (TIRF) microscopy. The laser illumination was focused at an angle of 54.5° with respect to the default position. Images of a field of view (FOV) measuring 80 × 49 µm and movies were taken with a laser intensity kept constant at 780 kW/cm^2^ for the green (532 nm) laser. After mounting the flow chamber onto the microscope, the sample was rinsed with 2x SSC. Nanodiamonds (90 nm, Cytodiagnostics, Canada) were diluted 1:10 in 2x SSC, added to the chamber and incubated until ~20 particles per FOV were visible. Subsequently, the sample was washed with DNA-PAINT buffer DB (5 mM Tris-HCl pH8, 75 mM MgCl_2_, 1 mM EDTA, 0.05% Tween-20) that had been described previously [55]. Before every imaging round, an imager targeting one of the gene segments or sequences was diluted in DB and the sample was imaged. After every round, the chamber was washed with DB for 2 min. This procedure was repeated until all sequences had been imaged for the desired number of times. Depending on the goal of the experiment – super-resolution microscopy or segment stoichiometry – the imager concentration, exposure time, total imaging time and the imaging mode were adjusted (Table S3). Moreover, to obtain the same average number of binding events for every segment, the imaging times were adjusted for every barcode-imager combination (Table S1).

### Image Processing

The DNA-PAINT software platform Picasso was used to obtain and process localizations from the ‘.tif’ files acquired by the Nanoimager [49, 56]. First, the raw data was cropped using imageJ to exclude an area with a lower illumination. The processed raw data was then opened in Picasso *Localize* and the localization algorithm was run with the following parameters: Box side length: 7, Min.net gradient: 3000, Baseline: 400, Sensitivity:2.75 Quantum efficiency: 0.82, Pixel size: 117, magnification factor: 0.65.

In addition to the x and y positions of the segments, we determined the axial positions of PAINT localizations using an astigmatic lens. We estimated the gain and baseline of our camera by the sCMOS analysis provided by GDSC ImageJ plugins [57]. The axial scaling factor is estimated using the GATTA-PAINT 3D HiRes 80R Expert Line (GattaQuant, Munich, Germany, **Fig S13**) [58]. We first detected each nano-rod by selecting a few examples and subsequently using the “Pick similar” function in Picasso render, and then determined the centre of the base and top of each nano-ruler using a Gaussian mixture model. The distance between two centres (the measured length of nano-ruler *L_m_*) and the measured angle between the nano-ruler and the surface *θ_m_* should follow the equation

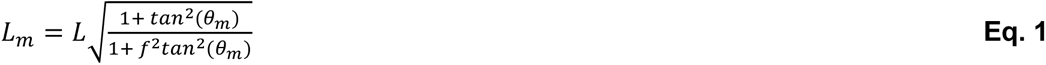

where f is the axial scaling factor and L is the actual length of the nano-ruler [58]. According to our fitting, the length of nano-ruler was ~77 nm, very close to the provided value of 80 nm. After localization, all resulting ‘.hdf5’ files were loaded into Picasso Render, the fiducial markers were visually identified, picked and used to undrift the localization file for each segment.

### vRNP structural analysis

After undrifting, the fiducial markers were used to align the localization files. By calculating the average squared distance of localizations to the centre of the fiducial marker to determine, we identified the fiducial markers which were immobilized during the recording. The shifts between two consecutive channels and their squared Mahalanobis distances were calculated. The fiducial markers with squared Mahalanobis distances larger than the quantile of 0.68 of a chi-distribution were considered as mobile during buffer exchange. After identifying the properly immobilized fiducial markers, the transformation matrix between two consecutive channels is calculated using static fiducial markers by point-to-point Iterative Closest Point (ICP) algorithm [59, 60]. The transformation matrices of subsequent channels are calculated simply by matrix multiplication.

After the localization files had been aligned, the localizations were linked to avoid bias due to dwell length (Max. distance: 1.00, Max.transient dark frames: 1.00). To identify segments, localizations were clustered using DBSCAN in Picasso Render. The DBSCAN parameters depended on the localization counts per segment and the localization background (default values: Radius = 0.5, min. density = 25). Once all gene segments in a FOV had been detected as described above, they were clustered together to complexes (referred to as “particles”) by agglomerative clustering (distance threshold = 234 nm). Once the particles had been indexed, the x and y positions of the localizations in each segment were used to fit the vRNP spine according to the following procedure: the outline of the segment was reconstructed by rendering every localization as a Gaussian (sigma = 3.9 nm, pixel size 0.234 nm). The resulting image was subjected to a low-pass order-one Butterworth filter with a cut-off frequency of 0.02 and Gaussian blur (sigma = 5.85 nm) and converted to a binary by thresholding with the ISODATA method [61]. A Hilditch skeleton was fitted to the binary image and all branches but the longest ones were eliminated to obtain the final vRNP spine, the length of which was considered to be the length of the vRNP (Fig 4E, center) [62]. The orientation of the segment in 2D was determined by fitting the skeleton with a straight line and calculating its angle with respect to the image axes (Fig 4E, right).

### Structural analysis of individual virus genomes

For the analysis of segment distances below, we considered particles of any shape, since filtering strategies based on the degree of vRNP alignment, on the assumed particle orientation, and on particle size did not increase the correlation between the experimental replicates. Analysis of vRNP distances within the virus particles demonstrated a moderate correlation of the lateral (x,y) and axial (z) directions (**Fig S14**), which was why we included the z positions in the calculation of the vRNP distances (see below). However, the average vRNP distances in z were only about one third of the x/y distances, suggesting that either the virus particles may have been slightly flattened during the preparation process, and/or that the fraction of elongated particles preferentially aligned with the surface.

### High-throughput stoichiometry analysis

The clusters were found by using a DBSCAN method for low-dimension Euclidean space with an eps = 1 and min_samples =4 [63]. The channels were aligned by redundant cross correlation (RCC) [64]. The positions of the clusters (average x and y values of the localizations) were saved, and fluorescent traces were extracted from the corresponding positions in the raw data. Binding and unbinding events in the traces were detected by a PELT analysis [65] and the traces were considered as “segments” if the number of binding events exceeded an adaptive threshold determined by analysing the binding event distribution of the observed segment population (Fig S1, Fig S3).

To control for the non-specific binding of both barcoded probes and imagers to the surface, the experiment was repeated with an oligonucleotide set that had the same number of sequences with the same length and the same barcode, but the DNAs of the set were non-complementary to NA vRNA (and to any other viral RNA) (red line, **Fig 1D**). Upon comparing the two distributions, we observed that only the population of traces with the high number of events corresponded to sequence-specific binding of barcoded probes (and therefore, to vRNPs), while the population with low number of events was due to background binding. Under the assumption that a Poisson distribution at the observed rate of events (λ ≈ 40) approximates a normal distribution, the population was fitted to determine a detection threshold (specifically, a number of binding events) where the probability of a false positive and a false negative NA vRNP detection was equal (**Fig S1**). For the remaining gene segment-imager pairs, the imaging time was adjusted so that the average number of binding events approximately corresponded to those of NA and P3 (n = 34 ± 4.6 STD, **Fig S2**, **Table S1**). In locations where traces occurred during multiple segment read-outs, the segment-specific population was more prominent in comparison to the background, increasing the odds of a true positive detection for a given event count **(Fig S4)**. Hence, the detection thresholds for every segment were calculated depending on how many other segment read-out rounds in the same position showed fluorescent traces.

**Figure 1.**
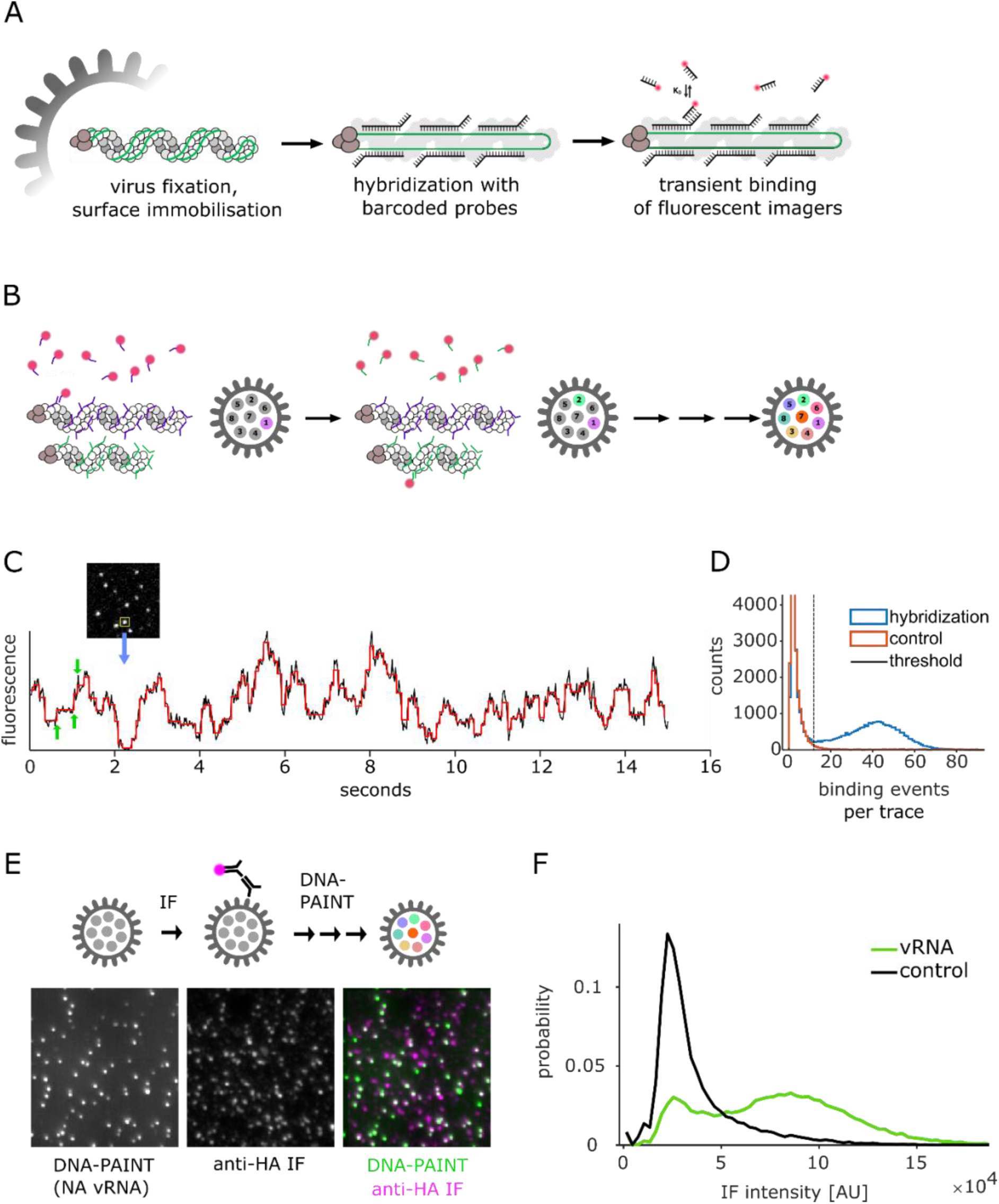
A DNA-PAINT protocol for the segment-specific detection of viral vRNPs. A) vRNP detection principle. Virus particles are surface-immobilized, fixed, permeabilized and incubated with non-fluorescent barcoded probes, that bind to 20-50 different sequences on a viral RNA segment. All probes carry the same segment-specific oligonucleotide extension that is transiently bound by fluorescent imager strands. B) Several combinations of barcode extensions and imager strands can be used to sequentially to detect various gene segments in a single virus particle C) Snapshot of a diffraction-limited movie of NA vRNPs inside viral particles incubated with a high concentration (50 nM) of complementary imager strands, and recorded fluorescence intensity time-trace of highlighted spot. Black: fluorescence signal, red: step detection via PELT analysis [65]. Step increases are interpreted as imager binding events. Green arrows: Three consecutive binding events. D) Distribution of binding events per spot when imaging NA vRNPs. Blue histogram: Assay with barcoded probes complementary to the NA segment. Red histogram: Negative control with non-complementary barcoded probes that carry the same barcode. Dashed vertical line: Threshold for the detection of viral segments. E) Experiment to demonstrate co-localization of vRNA and the viral envelope: Individual viral envelopes were stained via anti-HA IF, followed by image acquisition and high-throughput DNA PAINT (B-D) of the same positions. vRNA (left panel) and HA (middle panel) showed extensive co-localization (right panel). F) Distributions of HA IF intensity detected in positions with vRNA (green) and random positions in a control experiment (black).

## RESULTS

### Detection and identification of all 8 vRNPs inside single virus particles by DNA-PAINT

To identify and spatially resolve all eight gene segments inside single virus particles with nanometer precision, we developed a DNA-PAINT approach customized to detect viral particles (**Fig 1**).

Using 20-nt hybridization sites on the RNA segments, we designed *in situ* hybridization (ISH) probe sets for each of the eight viral segments of influenza strain PR8, ranging from 18 to 48 primary probes per segment, depending on gene length and site availability (**Fig 1A, Table S1**, Material and Methods). In our experiment, we immobilised purified and fixed PR8 viral particles on the coverslip by evaporating the virus solution. We then hybridized each viral RNA segment with its set of primary DNA probes (**Fig 1A**, middle panel); although the primary DNA probes were non-fluorescent, they carried 9- to 10-nt oligonucleotide extensions (“barcodes”), which were then bound transiently by fluorescent imager strands in a subsequent read-out step (**Fig 1A**, right panel). The resulting fluorescent signals were recorded by a TIRF microscope and used to detect and locate vRNPs inside the immobilized virus particles.

Our approach has several advantages **(Fig 1A)**. First, hybridization of several primary probes to one viral RNA segment (multiplexing) increases the signal of individual influenza vRNPs significantly above the background of unbound probes [11]. Second, the experiment was conducted in TIRF mode, where the sample is only illuminated within a thin section above the coverslip, thus providing a virtually unlimited reservoir of fluorescent imagers that are not illuminated (and thus not subject to photobleaching) prior to binding [66]; consequently, the signal-to-background ratio of detecting individual vRNPs can be optimized by increasing the sampling time. Third, by adjusting the imager concentration, individual hybridization events can be localized with very high spatial precision, leading to reconstructed super-resolution images of the target structure (PAINT). Fourth, since the imager binding is transient, the imagers can be swiftly exchanged under non-denaturing conditions, allowing the read out of several barcodes sequentially while preserving the structure of the immobilized virus particles **(Fig 1B)**.

As a proof-of-principle, we set out to detect neuraminidase (NA) vRNPs rapidly (20 s for a FOV with ~ 500 vRNPs) by incubating the immobilised particles with barcoded probes hybridized in 50 nM of the complementary imager P3 **(Table S3)**. At this concentration, several imagers bind to a single vRNP at the same time, typically resulting in bright fluorescent point sources (“spots”) with a stepwise increase and decrease in fluorescence over time (trace), as imagers were binding and unbinding, or - to a lower extent - bleaching **(Fig 1C)**. The resulting frequency distribution of binding events was compared to a control probe set that was unable to specifically bind imagers, which enabled us to detect segments by filtering out traces with an insufficient number of binding events (**Fig 1D, Materials and Methods**). A comparable signal could be detected for the remaining 7 segments (**Fig S2**). Hence, we demonstrated that we could fluorescently label and detect individual vRNPs in situ in a selective, reversible manner and a mild chemical treatment.

To ascertain that we were observing whole virus particles (and not groups of segments dissociated from viral particles), we applied an anti-haemagglutinin (HA) immunofluorescence (IF) co-staining to image the exterior of the virus particle **(Fig S3)**. Subsequently, we performed our high-throughput DNA-PAINT assay, followed by labelling and imaging of same positions in the sample **(Fig 1E)**. The apparent co-localization of the accumulated DNA-PAINT fluorescence (i.e., segments) with IF spots (i.e., glycoproteins embedded within the viral envelope) was high. We then measured the HA IF in a diffraction-limited spot around particle positions indicated by DNA PAINT, and compared their distribution with the IF signal from random positions in the sample. We determined that 73% of particles had an IF signal above the conservatively estimated background (5 x 10^4^ A.U.) and thus carried HA, confirming that we were primarily observing complete virus particles **(Fig 1F)**.

### High-throughput analysis of segment content in virus particles points to a selective and efficient genome assembly

For a sequential interrogation of all vRNPs in a virus sample, we typically detected and localized ~17,000 vRNPs per read-out round. For technical replicates, we shuffled the segment read-out sequence to control for potential changes in the vRNP detection sensitivity over the course of the experiment. We found that the majority of segments formed clusters with a distance < 234 nm (2 pixels) from each other, in agreement with previous FISH studies on influenza samples [11, 47]; we will thus refer to those clusters as “virus particles” throughout this study. We also excluded the presence of more than one gene for both PA and HA in the vast majority of particles (~ 94%), in agreement with Chou et al. [11] **(Fig S5)**. The average probability of co-localization between two segments in our study was 74% (± 4% STD).

When we statistically analysed the number of segments per particle, we found that most particles (82% ± 5% STD) did not contain a full set of segments and could therefore not cause a productive infection by themselves **(Fig 2A)**. The number of gene segments within particles ranged from 1 to 8, with particles having either just one or 7-8 segments being overrepresented in the population compared to particles with an intermediate segment count (2-6). The described distribution of segments resulted in a higher fraction of particles with the full set of 8 segments than expected by a stochastic combination of segments with the same co-localization rate, i.e., when we stochastically distributed the segments across complexes, segments with intermediate segment counts were more abundant (**Fig 2A**, white bars). We further characterized the assembly efficiency at a segment count N by the ratio

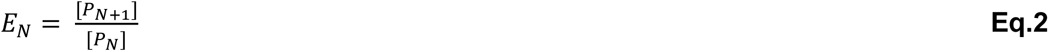

where [P_N+1_] and [P_N_] are the concentrations of the ternary complexes with the respective segment counts, we found that the assembly efficiency increased by ~ 0.25 per added segment and peaked at 1.5 (P_5_), reminiscent of a cooperative process **(Fig S6)**. When the last segment had to be added (P_7_), the assembly rate dropped off again.

**Figure 2.**
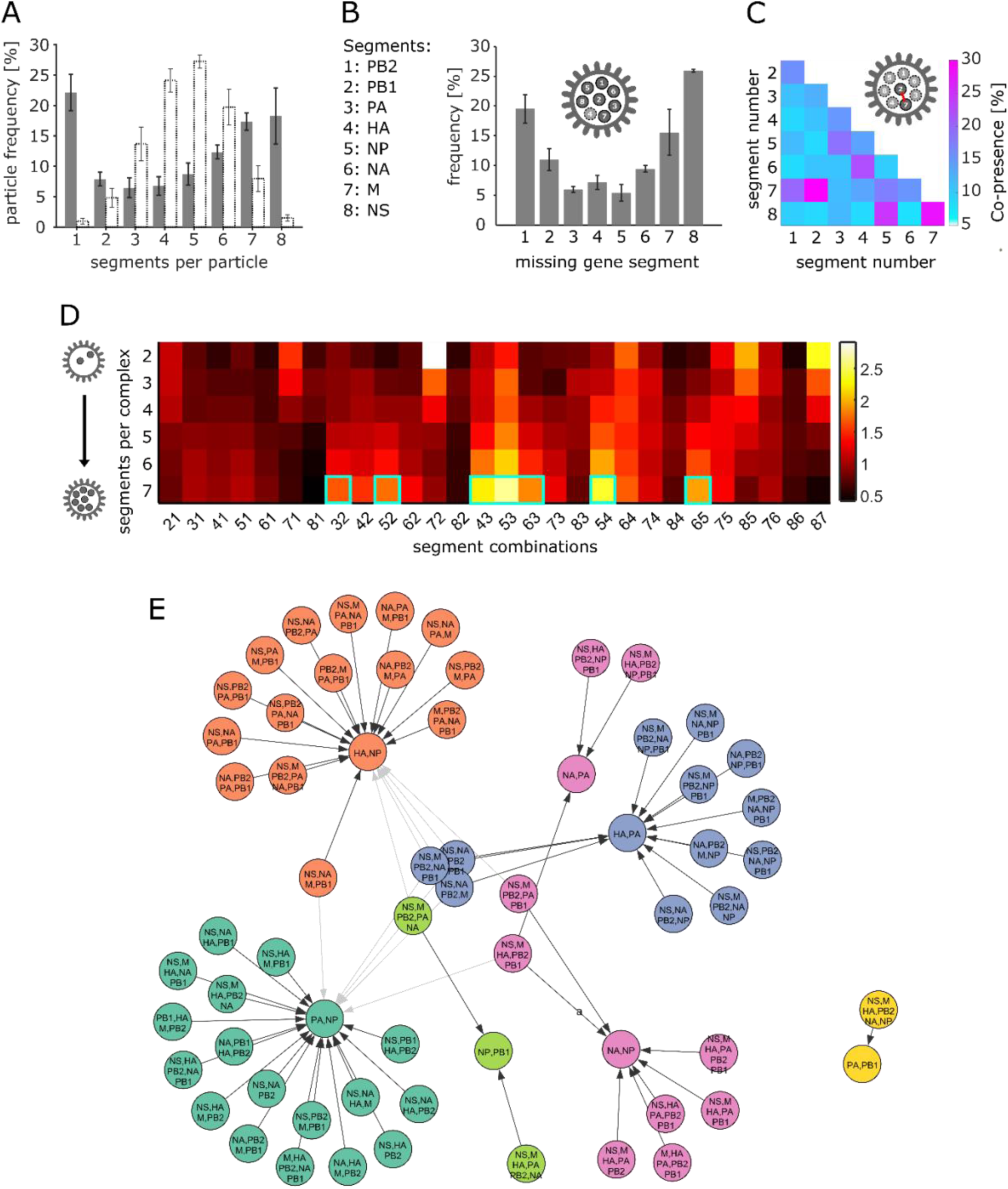
High-throughput analysis of segment stoichiometry and co-presence in individual virus particles. Error bars: standard deviation (SD) of three technical replicates. A) Number of individual gene segments per particle (grey bars), and comparison with same dataset with segments shuffled between particles to simulate a stochastic distribution (white bars). B) Frequency of gene segments that are absent from particles with only one segment missing. Legend: Segment numbers that code for genes, used in B-D. C) Segment co-presence in particles with two segments for every possible combination. Co-presence of two segments is defined as the fraction of the number segment pairs and the average total number of both segments. D) Normalized segment co-presence (Eq.3) in particles with higher segment counts. The first horizontal line represents the data shown in F. Cyan boxes: Segment pairs that form association centers in the FP analysis (see below). E) Network of prominent associations between vRNP subcomplexes obtained via a frequent pattern (FP) algorithm ([67], Figure S8). The resulting association rules were arranged into a network and communities (colors) were identified via the Leiden Algorithm [70].

We then examined the role of specific segments during genome assembly. One possible interpretation of the 7+1 pattern observed in budding virus particles is that it arises from the organization of all segments around a particular, central segment that makes several contacts to all other segments; such an organisation suggests that the central segment is likely to play a crucial role for genome assembly and is very unlikely to be absent from 7-segment particles. To test this hypothesis, we analysed all particles containing 7 segments and found that no segment stood out in such a way, although the relative absence ranged substantially (from 5% for the NP segment to 25% for the NS segment), indicating varying degrees of involvement for individual segments in selective packaging **(Fig 2B)**.

In short, we assessed the RNA segment content of a large number of virus particles in purified PR8 virions and found that the majority of them did not contain a full set of genes. However, the fraction of particles with a complete genome was higher than expected from a purely stochastic distribution of the segment counts among the particles, which can be explained by an increased efficiency of genome assembly in the later stages, indicating the presence of cooperativity during assembly.

### Quantification of segment interactivity based on the co-presence of segment pairs in the virus population

We then turned our attention to the interaction between individual gene segments. We based our approach on the assumption that an RNA-RNA interaction during selective packaging would manifest in a higher probability of both segments being present (co-presence) in particles with a partially assembled genome (i.e., assembly intermediate on the path to fully assembled viral particles).

We first examined the co-presence in two-segment particles (P_2_), as we expected more statistical noise from the stochastic co-presence in particles with more than two segments **(Fig 2C)**. When we compared the co-presence of the segment pairs between three technical replicates, we obtained Pearson coefficients (PC) in the range of 0.89-0.95, underlining the reproducibility of the analysis **(Fig S7)**. The individual co-presence probabilities of all segment pairs ranged from 6 to 30% for two-segment particles (**Fig 2C**), with a relatively low number having a particularly high co-presence probability (e.g., PB1/M, NS/M, and NP/NS), and all remaining segment pairs forming a fairly uniform plateau of low co-presence probabilities. This result pointed towards a complex network of a multitude of RNA-RNA contacts, as suggested previously [31]. However, certain stronger contacts might play a more important role in selective packaging. Interestingly, the segment pairs with the highest co-presence involved at least one of the shortest segments (M and NS).

After verifying that the experimental replicates showed a high correlation (average 0.8 – 0.93) for all incomplete multi-segment particles (**Fig S7**), we normalized the co-presence probability with the following equation:

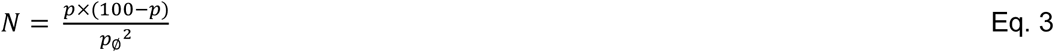

where p is the co-presence probability of the segment pair in particles with a specific number of segments [%] and p_ø_ is the average co-presence probability of all particles with the given segment count. The equation accounts for the fact that the co-presence naturally increases in particles with a higher segment count, and the fact that the deviation of the co-presence from 100% is key for characterizing segment pairs with increasing co-presence values. With the values obtained from this normalization, we plotted the co-presence for each segment pair depending on the number of segments in a particle **(Fig 2D)**. The relative co-presences for most pairs changed with the segment count, either gradually increasing or decreasing, which suggested that contacts mediating segment pairing can selectively contribute to earlier or later stages of the packaging process. A possible mechanism for how contacts gain importance towards the later stages of genome assembly is their mutual stabilization in multi-segment intermediates. This can also explain the apparent co-operative packaging behaviour (see above). The highest co-presences for 7-segment particles involved pairs of segments that have been identified as being rarely absent from particles (PA, HA, NP; see **Fig 2B**), underlining their importance in the late assembly stages.

So far, our observations pointed to a complex relationship involving many RNA-RNA contacts between multiple segments with synergistic, cooperative effects. Hence, we identified the most significant associations between all of the 247 possible subcomplexes by association rule mining, where each gene segment was treated as an item and frequent item-sets (i.e., subcomplexes) in the observed transactions (virus particles) were identified using a Frequent Pattern (FP) Growth algorithm and subsequent filtering (using the parameters Lift, confidence and Zhangs metric [67–69]) (**Fig S8**). After filtering, the remaining 67 association rules formed six communities, which we visualized via the Leiden algorithm (**Fig 2E**, [70]). Each of the communities appeared to have a core structure that was involved in a disproportionate number of associations. Intriguingly, these 7 subcomplexes each contain two segments, which also represent the pairs with the highest co-presence in 7-segment particles (see 7 cyan boxes in **Fig 2D**), illustrating their key role in later stages of genome assembly.

To assess whether genome assembly in other influenza A strains follows comparable rules, we ranked all combinations with a specific segment count by frequency, as has previously been done for intracellular vRNP subcomplexes in H3N2 (A/Panama/2007/99)-infected cells (**Fig S9** and Haralampiev et al., 2020, Fig 4A, [47]). We compared mature, budded virus particles (PR8) with intracellular virus progenitors (A/Panama), as the numbers of mature A/Panama particles in Haralampiev et al. were insufficient to perform the same type of analysis. A qualitative comparison of the plots showed that the distributions of stoichiometries in both datasets had similar shapes, tentatively suggesting that the molecular interactions shaping both genomes are comparable in strength and number. However the most frequent combinations in A/Panama were separated from the others more clearly and could be interpreted as parts of preferential assembly pathways, which could not be deduced from our data on PR8. Apart from a few notable exeptions (127, 1278), the identitities of frequent combinations in both virus samples differ, suggesting that the genetic variation between the two subtypes generally leads to different interaction motifs and assembly routes.

In short, we quantified the probability of both segments residing together in a virus particle (co-presence) in order to assess the interactivity of all segment pairs in our virus sample. We found that, for various pairs, the co-presence values changed dramatically with an increasing particle-segment count, an indication that the early and late stages of genome assembly are directed by different interactions.

### Structures of vRNPs inside virus particles visualized by DNA-PAINT

We then turned our focus towards spatially analysing the genome structure of virus particles with DNA-PAINT by reducing the concentration of the imagers for each segment to 5 nM and increasing the imaging time by a factor of 30 (**Table S3**). In contrast to the data gathered in the high-throughput experiment (**Figs. 1-2**), the binding events were now spatially separated and resulted in clusters with an average of ~100 localizations for the NA segment **(Fig 3A, B)**. If the RNA-binding sections of the barcoded probes were exchanged by non-binding oligonucleotides, localization clusters of a similar size were a lot less frequent, confirming the specific binding of both barcoded probes and imager strands.

**Figure 3.**
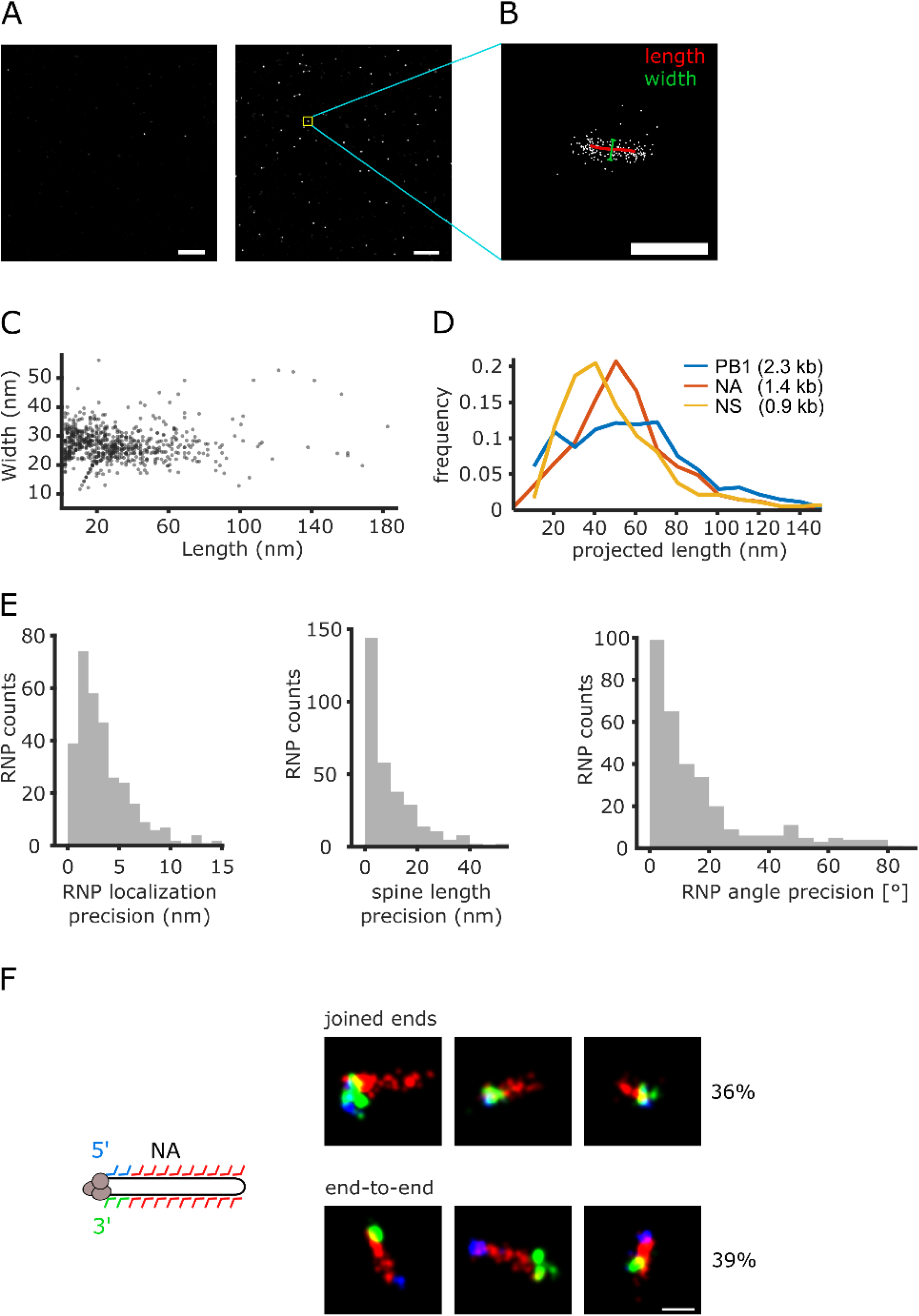
Characteristics of vRNPs visualized by DNA-PAINT inside immobilized virus particles. A) Overview of several super-resolved vRNP clusters. Left: Negative control with non-binding barcoded probes. Right Complementary barcoded probes. Yellow box: vRNP cluster sown in B. B) Typical localization point cloud depicting a super resolution image of an NA vRNP, with the skeletonized outline depicting its estimated length in x and y (red) and a bar depicting the width (green). Scale bar: 100 nm. C) Scatter plot of NA vRNP widths and lengths in a DNA-PAINT experiment. Every data point depicts the length to width ratio of an individual vRNP. D) Distribution of vRNP lengths from lysed virus particles for viral segments PB1, NA and NS (blue, red and yellow). E) To obtain the precision of determining the vRNP XY localization (top left), spine length (top right) and XY orientation (bottom), the localizations making up a NA vRNP point cloud were randomly split into two equally sized point clouds and the differences of the aforementioned properties were plotted as distributions. F) To investigate the quaternary structure of NA vRNPs after preparation, barcoded probes close to the 5’ and 3’ end contained additional barcodes (blue, green) to visualize the ends in separate imaging rounds. In 75% of the observed cases, both ends were found on either end of the outline (red), either on the same end (top row) or opposite ends (bot tom row). Scale bar: 50 nm.

We fitted a spine to the long axis of these clusters (***Materials and Methods***) to determine their length and width, and found a broad length distribution, ranging from 0 to ~90 nm **(Fig 3C)**. This range was consistent with the previously estimated length for NA vRNPs and the expectation that the particles and vRNPs inside them assumed random orientations during immobilization. In contrast, the average width was 25 nm and broadly independent of the measured length, in agreement with vRNPs of a previously estimated width of 25 - 30 nm [2]. The apparent lengths of vRNPs from lysed virus particles (pre-treated with 0.5% Triton X-100 and sonicated) increased with the gene length (NS = 0.9 kb, NA = 1.4 kb, PB1 = 2.3 kb), serving as an additional confirmation that vRNP outlines could be successfully resolved **(Fig 3D)**.

To understand the limitations of our imaging procedure, we determined the precision of the position, the length, and the axis orientation of the imaged vRNPs by randomly splitting the localizations making up a vRNP in half and determining the difference for each parameter between the subclusters **(Fig 3E)**. With this approach, we determined median precisions of 4 nm, 10 nm and 10° for the vRNP position, i.e., the mean position in x and y of all localizations that were part of the vRNP cluster, as well as the length and axis orientation (calculated from the vRNP spine, see ***Materials and Methods***) respectively, and concluded that determining these parameters for all segments inside a particle could be realized experimentally.

To assess the conformation and structural integrity of the vRNPs, we exploited the option to add different barcodes to individual primary probes and sequentially applied different imager strands to image the outline, the 5’ terminus, and the 3’ terminus of NA vRNPs inside immobilized particles **(Fig 3F)**. Out of 432 vRNP outlines, 357 (~83%) did not show signal on either one or both termini, possibly because the termini are inaccessible due to the interaction with the viral polymerase [71], or were not elongated enough to localize the termini to one end, possibly due to the stochastic nature of the deposition of the vRNPs on the glass surface. The remaining 75 showed sufficient signal for both the 5’ and 3’ terminus (> 4 localizations, all within a radius of ~10 nm of their mean position) and had an outline that was elongated enough to assess the localization of the ends (~40 nm or more). In 75% of these vRNPs, both termini were situated on the ends of the outline, with an almost equal probability of the termini being on either the same or opposite ends (36% and 39%), while the remaining 25% showed no clear localization of at least one terminus to either vRNP end.

Structural evidence in the literature strongly suggests that in intact vRNPs, both RNA termini are bound to the viral polymerase and hence situated on the same end of a vRNP [71, 72]. This vRNP conformation likely also applies to budded virus particles, where the polymerase was also found to be preferentially localized to the ends of vRNPs [73]. In our experiments, we found that the vRNP structures with separate termini were about twice the length of vRNPs with joined termini (62 vs 34 nm), suggesting that at least one of the RNA termini had detached from the polymerase and unwound from the vRNP. This suggests that we have a small number of partially denatured vRNPs in our experiment (~ 6% of the total number of vRNPs observed).

### Spatial analysis of the influenza genome with DNA-PAINT

Having shown that a large number of individual vRNPs can be structurally preserved inside an imaged particle, we set out to create super-resolution images of the viral genome by DNA-PAINT. To confirm that individual vRNPs inside particles kept their position throughout the experiment, we imaged NA vRNPs twice, first at the beginning, and later at the end of the entire imaging sequence **(Fig S10).** The median of the position differences before and after the imaging cycle was 15.6 nm, which is only a small fraction of the virion diameter (~ 100 nm), indicating that vRNPs generally stayed in the same location inside an immobilized particle, and opening the way for studying the genome structure by statistical analysis of a large set of particles.

Individual particles showed vRNP arrangements in various shapes, as illustrated in **Fig 4A** and **Fig S11**. While some particles were of a globular shape and contained vRNPs that appeared short and barely intersected (**Fig 4A**, top row), others consisted of mostly elongated, aligned vRNPs (**Fig 4A**, middle row). These observations are consistent with EM studies that show elongated particles with aligned genome contents, either in an axial or lateral orientation [39, 40]. In contrast, other virus particles had a more irregular vRNP arrangement (**Fig 4A**, bottom row); such an observation was also made for virus populations in EM data.

**Figure 4.**
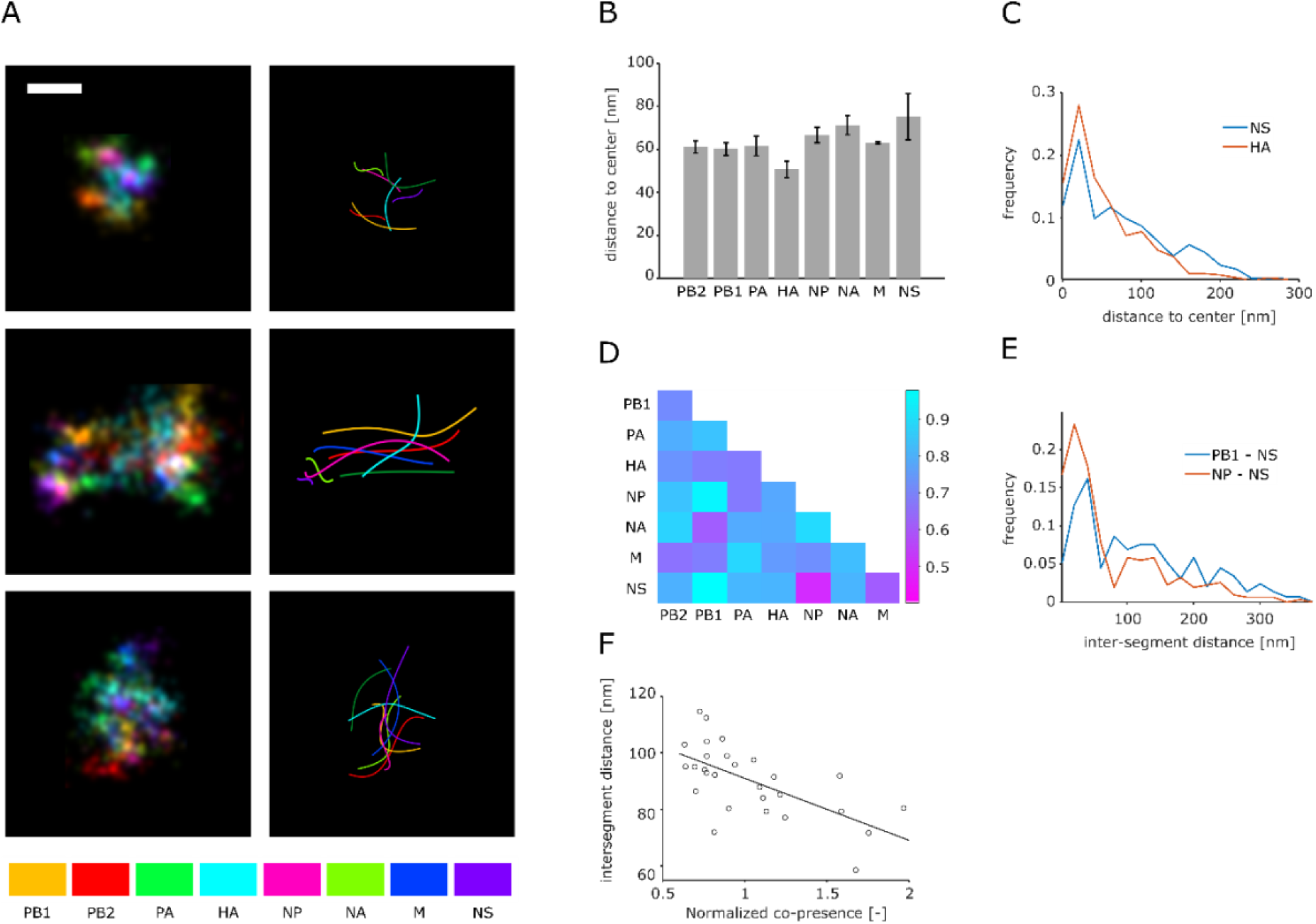
Spatial analysis fo the influenza genome with DNA PAINT. A) Super resolution images of the influenza genome. Different colors mark different segments within the particles. The examples include particles with clearly separated vRNPs in the XY (top row), vRNPs that are aligned and projected in XY (center row), and disordered vRNPs (bottom row). Left column: Images reconstructed from localizations and a gaussian blur of 3 nm. Right column: Spines of the detected vRNPs. Scale bar: 50 nm. B) Median segment to center distance of all segments. Error bars: Standard deviation after normalization by the average distance for all segments of three technical replicates. C) Distribution of segment-to-center distances for the shortest (HA) and longest (M) measured distance. D) Median inter-segment distances of three technical replicates. The color indicates the distance according to the bar on the right. E) Distribution of inter-segment distances for the closest (NA - NS) and furthest (PB1 - NS) segment pair. F) The inter-segment distance for every segment pair is plotted against its co-presence in particles with 2-4 segments. The correlation coefficient is given. Line: Linear fit of the data set.

Moreover, some segment aggregates that were recognized as particles by our analysis covered a larger area than that expected from a single virus particle. While this may suggest a partial disruption of the genome, large membranous structures with spatially separated genetic material were also observed in EM studies using the same virus preparation [40]. We observed that the transition between these genome appearances in our data was continuous, pointing to the entire particle population having a continuum of orientations with respect to the surface and a varying degree of structural organization.

We reasoned that increasingly assembled genomes would appear more structured; we thus exclusively analysed distances in particles with more than 6 segments. First, we examined the distances of each segment to the particle center to identify segments that would potentially form the core of a 7+1 configuration **(Fig 4B)**. The median distances of all segments to the center (~ 60 nm) were considerably larger than expected from EM data of budding virus particles, which reflects different particle orientations, as well as potential structural disruption by virus maturation and sample preparation, and the flattening demonstrated above (Materials and Methods). Likewise, both the closest and furthest segments (HA and M) had a broad distance distribution with an approximate spread of 150 nm **(Fig 4C)**. Even though the average median distance between any two segments varied between technical replicates (46 – 92 nm), the correlation of median inter-segment distances between the replicates (PC = 0.75 – 0.81) was sufficient to conclude that the genomes in our virus populations were structured to some extent (**Fig S12**).

To illustrate the significant difference in center-distance between the segments, we normalized the distances in each replicate to the average value of all segment pairs in all replicates (63 nm) and calculated the standard deviations from the normalized data (error bars in **Fig 4B**). Interestingly, the four longest segments (PB1, PB2, PA, HA) have a shorter center-distance than the remaining ones, in agreement with EM data from budding virions, where one of the long segments was always found in the center of a 7+1 configuration [38]. Interestingly, the segment with the shortest center-distance, HA, was also found in two of the three 2-segment subcomplexes with the most association rules in the FP analysis (**Fig 2E**) and might thus have a central role in organizing the genome during the packaging process.

### Mapping intersegment distances reveals that segment interactions shape the vRNP configuration inside mature virions

We also determined all 28 median inter-segment distances in analogy to the co-presence analysis (**Fig 2C, 2E**). Reminiscent of the center-segment distances discussed above, we measured a sizeable spread of the length distribution for segment pairs both close and apart (~ 250 nm) and a considerable variation of the average distance between replicates, once again indicating less tight associations than in budding virus particles **(Fig 4D-E)**. Nevertheless, the correlation of inter-segment distances across all replicates was high (PC = 0.74 - 0.81), suggesting an inherent structure of the genomes with median intersegment distances ranging from 41 – 67 nm.

We then asked the question whether the genome arrangement was caused by the inter-segment interactions that our co-presence experiments suggested. Indeed, we found a moderate negative correlation (PC = −0.61) between the observed inter-segment distances and the co-presence in 2-, 3- and 4-segment particles, indicating that interactions of two segments led to shorter distances between them **(Fig 4F)**. The correlation with inter-segment distances rapidly dropped for co-presence values of > 4-segment particles (**Fig S15**). A possible explanation is that these interactions, albeit important, might be weaker on their own and only become important in concert with other interactions as suggested above (**Fig 2D-E**).

We conclude that we were able to directly identify and visualize all eight gene segments inside influenza particles for the first time. We found that their configurations inside the particles show a pleiomorphy reminiscent of unidentified vRNPs in EM data. However, as suggested by locating the vRNA termini, the particles may be subject to some degree of internal disruption (see above). Analyzing segment distances from the particle center revealed a tendency for longer segments to occupy the particle center, recapitulating EM studies, while the inter-segment distances partially correlate with segment interactivity suggested by our stoichiometry analysis. These results support the notion that despite being segmented, the gene content of influenza particles remains structured after budding.

## DISCUSSION

In this study, we used DNA-PAINT to quantify the interactions of individual segments in influenza PR8 by quantifying packaging defects and assessing the spatial arrangement of vRNPs in many individual virus particles.

### Insights into influenza genome assembly mechanisms

Firstly, we detected that *any* segment can be the one missing from otherwise complete particles. This observation strongly suggests that no single segment is absolutely essential for an efficient packaging process, albeit some segments are missing more frequently **(Fig 2B)**. These observations differ from those in other segmented RNA viruses like Bluetongue virus, where a core of short gene segments is essential for the recruitment of the longer segments and is therefore unlikely to be missing from near-complete genomes [74]. In line with this, we detected every possible combination of segments, which points towards a complex interaction network, as suggested previously [31]. Nevertheless, a small number of segment pairings were detected at an elevated frequency, pointing towards sequence-specific interactions between them **(Fig 2C, D)**.

Notably, the interactivity of segment pairs seems to change with the number of segments per particle, with some segment pairs increasing in interactivity **(Fig 2D)**. Hence, if we interpret particles with an incomplete set of segments as intermediate stages on the assembly pathway to a full genome, our results suggest that the later assembly stages are dominated by interactions that are enhanced by the presence of additional vRNA segments. This property is likely to rely on the synergistic action of RNA contacts between multiple segment pairs, which is consistent with the fact that multiple segment pairs seem to be involved in selective packaging **(Fig 5)**. Moreover, synergy between RNA contacts is supported by the observed efficiency increase in later stages of virus assembly, which indicates cooperativity between multiple segments **(Fig S5)**. Recent observations that viral RNA structures are more open in virio and in vivo suggests another plausible mechanism for the shift in segment interactivities during assembly: As suggested for mature viral particles, the RNA in larger subcomplexes may be less structured due to crowding effects and show reduced binding of host proteins, facilitating the formation of additional RNA contacts between vRNPs [34, 75].

**Figure 5.**
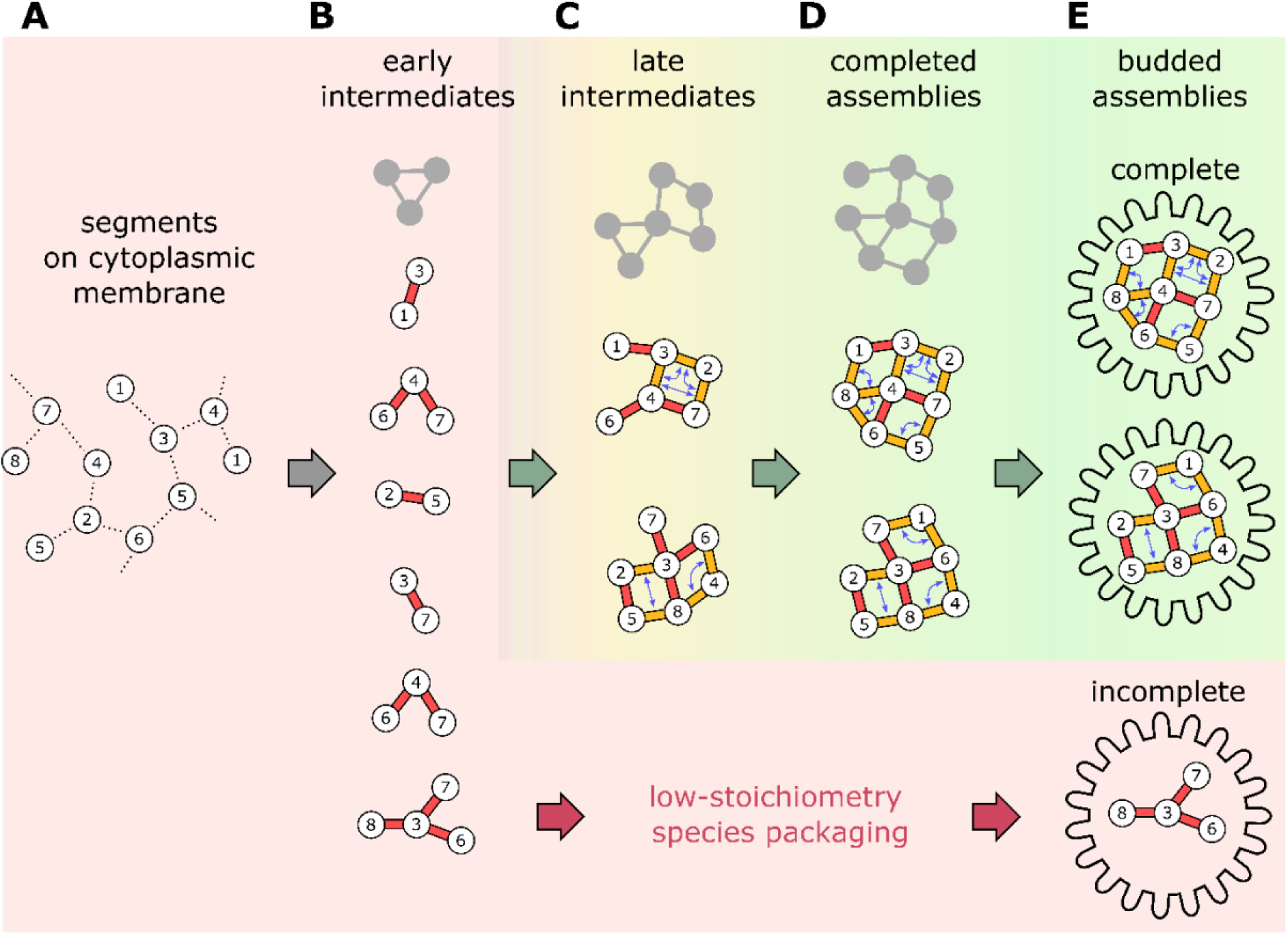
A model for influenza genome assembly via multiple pathways involving cooperativity between RNA-RNA contacts. A) Newly synthesized vRNPs are concentrated at a membrane (e.g. during vesicular transport to the cytoplasmic membrane. [9]) Unspecific interactions (dashed lines) between all segments locally concentrate the vRNPs to later facilitate segment-specific interactions. B) In the early stages of assembly, strong interactions (red sticks) between individual segment pairs form low-order subcomplexes. C) In the later stages, weaker interactions (orange sticks) co-operatively stabilize each other (double arrows) to form higher order complexes (green arrows), resulting in the assembled genome (D), which is packaged into budding virus particles (E). The illustrated assembly routes merely represent conceptual examples, as a multitude of RNA contacts results in many alterative assembly routes (grey outlines). If lower order subcomplexes take too long to grow to sizes at which cooperative effects accelerate assembly, incomplete viral genomes will be packaged (red arrows).

Overall, our results exclude the presence of a single pathway for influenza genome packaging. Further, our results exclude the presence of a completely stochastic combination of segments. Instead, our results strongly suggest the presence of several packaging paths that are outlined by sets of preferred interactions (**Fig. 5**). Such a kinetic landscape, which is conceptually similar to protein folding of moderate-side proteins, allows packaging to proceed both with high speed and efficiency.

Intriguingly, in the early stages of assembly, the highest interactivity is seen in pairs involving one or both short segments (M and NS) **(Fig 2D)**; such low-stoichiometry particles are likely to be viral assemblies that fail to grow sufficiently fast towards complete assemblies before budding (**Fig. 5**, red path). This suggests that the incorporation of M and NS into the viral particle depends on very few interactions that do not act in synergy with others. This property can be explained by their short sequence that might put tight constraints on the number of sequence-specific contacts they can make, and hence, on the number of other segments they can effectively interact with. Moreover, these two segments (M and NS) are among the three most frequently missing from otherwise intact particles **(Fig 2B)**. In agreement with our observations, mutations in only a few nucleotides of segment 7 (M) have been shown to drastically lower the number of infective particles in a virus population [15]. Taken together, these results suggest that the incorporation of the small segments (M, NS) is mediated by relatively few RNA-RNA contacts and is thus prone to failure. This potential “Achilles heel” in the formation of infective virus particles could be exploited by sequence-specific targeting of a few short RNA motifs in hypothetical influenza therapeutics.

In contrast, the interactions dominating the later stages of assembly appear to be predominantly between longer segments (PB1, PA, HA, NP, NA) with certain pairings being central in the network of subcomplex dependencies created by the FP analysis **(Fig 2E)**. Of these, the segment pairing with the highest co-presence value in 7-segment particles, segment 3 and 5 (PA, NP), were predicted to interact by an earlier study on selective packaging in PR8 [22]. If each of these segments interacts with multiple other segments as we reasoned above, this will result in a spatial genome organization, where at least some of these segments occupy a more central position. In fact, published EM tomographs show that the central position in the 7+1 configuration during virus budding is always occupied by one of the 4 longest segments [38]. Further studies on other strains may show whether the observed interplay of long and short segments is a general property of selective packaging in influenza.

### Structural integrity of virus particles imaged with DNA-PAINT

Preparing the virus sample for DNA-PAINT required various steps that may have altered the native structure of individual virus particles. We immobilized virus particles by evaporation, which turned out to be the most efficient way to reach a sufficient virion density for high-throughput studies while keeping background fluorescence low. However, despite influenza particles being able to survive desiccation, drying might deform the particle and introduce strain on its contents [76]. Moreover, the permeabilization agents and chaotropic ingredients of the hybridization buffer, as well as the oligonucleotide probes themselves, may cause some denaturation. Consistent with this, ~50% of vRNP structures where we could clearly observe both RNA termini appeared to have an open conformation, which has not been reported by previous structural studies.

On the other hand, the majority of vRNPs (>80%) had no signal for both termini, which suggests that vRNP denaturation and successful labelling of the termini might be correlated, resulting in a large over-estimation of the denatured fraction. Moreover, most particles appeared to retain their envelope as demonstrated by the presence of HA IF, suggesting the overall structural preservation of the virions. Future implementations of our method may further assess and further minimise sample denaturation.

### Insights into the influenza genome organisation

We found a wide variation within all segment-center distances and the segment-to-segment distances in our experiment, indicative of a flexible arrangement **(Fig 4C, E)**, with the caveat that the genome structure inside a purified influenza particles might be altered after budding, and/or during virus purification and preparation for imaging. However, the average distances measured for all segment-pairs and segment-to-center distances showed clearly reproducible variations, supporting a non-random arrangement that persists after the budding process.

Moreover, there was a moderate correlation between segment-to-segment distances in 6- to 8-segment particles, and the putative interactions in the early stages of the packaging process, as determined by the co-presences in 2- to 4-segment particles **(Fig 4F)**. The correlation between segment stoichiometry and genome structure was particularly striking, as they were obtained in an independent fashion with non-overlapping populations of the sample regarding the segment count per particle (see above). It is unclear, however, why this correlation does not hold up for putative interactions in the later stages. One possible explanation is the configurational flexibility of the potentially synergistic interaction network. Moreover, it is conceivable that the genome structure during assembly is indeed dynamic, with vRNPs swapping places during the process. Mechanistically, this could be explained by steric clashes between the relatively rigid vRNPs disrupting some of the weak late stage interactions. Lastly, weaker RNA contacts may dissociate due to denaturation happening between virus budding and imaging.

In agreement with the aforementioned EM data, one of the four longest segments, HA, had the closest median distance to the particle center in our DNA-PAINT experiments **(Fig 4B)** [38], making it the prime candidate for the center piece of a hypothetical rigid 7+1 configuration, where the central segment and potentially others have their fixed positions.

In our work, high-throughput PAINT-based multiplexing and nanoscopy proved to be an efficient tool to gather structural information on the influenza genome. While sequencing-based methods like SHAPE, SPLASH, 2CIMPL and others identify potential RNA-RNA contacts at the nucleotide level, their individual contributions to selective packaging can be evaluated through comparison with our experimental data that provide another puzzle piece to describe the exact packaging mechanism by documenting its result. Likewise, high-throughput, multiplexed nanoscopy is an efficient tool to characterize the architecture of biomolecular assemblies that are dynamic, flexibly arranged, consist of numerous individual components and therefore, cannot be reconstructed by particle averaging. As this methodology can be applied using any automated TIRF microscope with single-molecule sensitivity, and is still being developed and refined, we anticipate that it will become increasingly important for structural biology in all domains of life.

## ACKNOWLEDGMENTS

The authors thank the lab of Andrew Turberfield (Oxford Physics) for use of an ONI Nanoimager, all members of the Fodor lab for helpful discussions, and the MICRON Advanced Bioimaging Facility at Oxford (supported by Wellcome Strategic Awards 091911/B/10/Z and 107457/Z/15/Z) for access to an ONI Nanoimager in their facilities.

## AUTHOR CONTRIBUTIONS

C.H., N.R., and A.N.K. conceived the project. C.H., N.R. and A.N.K. designed experiments. C.H. and Q.Z. performed experiments and analysed data. Q.Z. performed simulations and provided software. C.H and Q.Z. analysed data. C.H., Q.Z., A.N.K., E.F. and N.R. discussed and interpreted data. C.H. wrote the paper, and all authors had the chance to read and edit the paper.

## DATA AVAILABILITY

Raw movies and images as well as localisation files for single molecules are available upon request.

