## Supplementary Information for "High-throughput Single-Virion DNA-PAINT Reveals Structural Diversity, Cooperativity and Flexibility during Selective Packaging in Influenza"

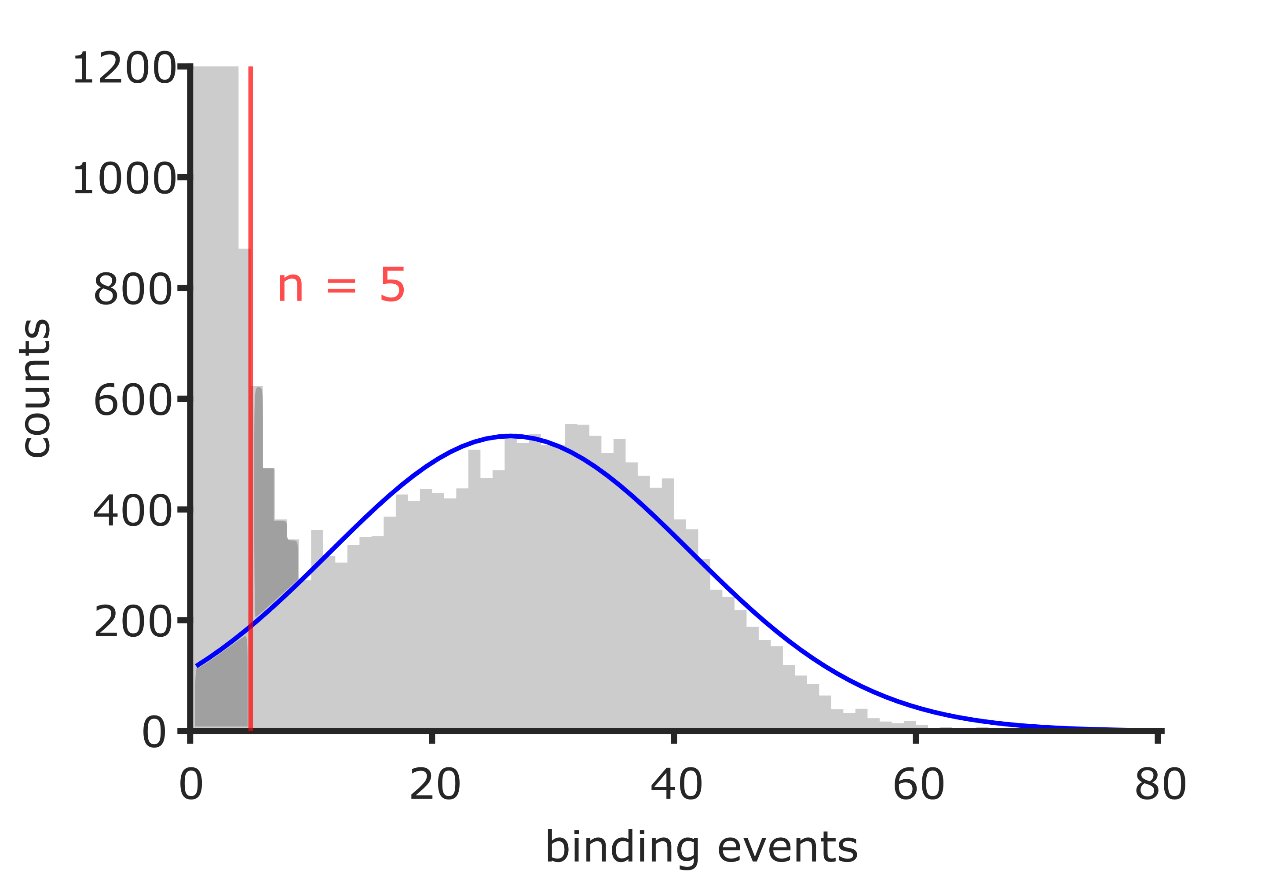

**Figure S1. Obtaining the threshold of binding events for detecting a vRNP** (segment NA, see also Fig 1D). The second part of the binding event distribution for fluorescent traces is fitted by a normal distribution (blue line). The number of binding events were the area above the curve to the right (false positives, dark grey) is equal to the area below the curve to the left (false negatives, dark grey) is n = 5 (red line), hence the threshold for vRNP is 6 binding events.

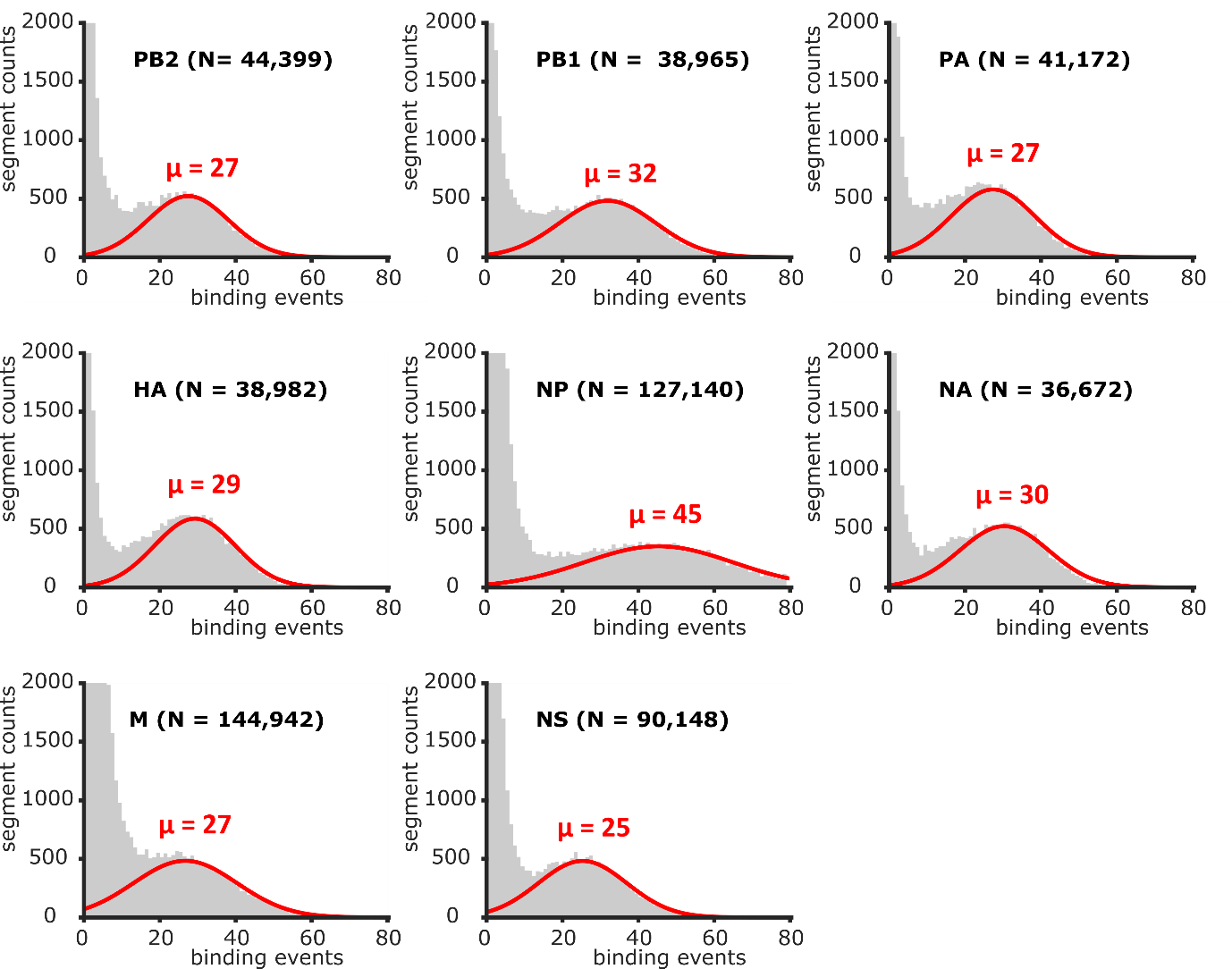

**Fig S2. Binding event distributions of spots with a persistent or recurring fluorescent signal from all 8 imaging cycles** (segments 1-8). Red line: Gaussian fit to the population corresponding to vRNPs (see Figure 1D). µ: Mean number of imager binding events to a vRNP

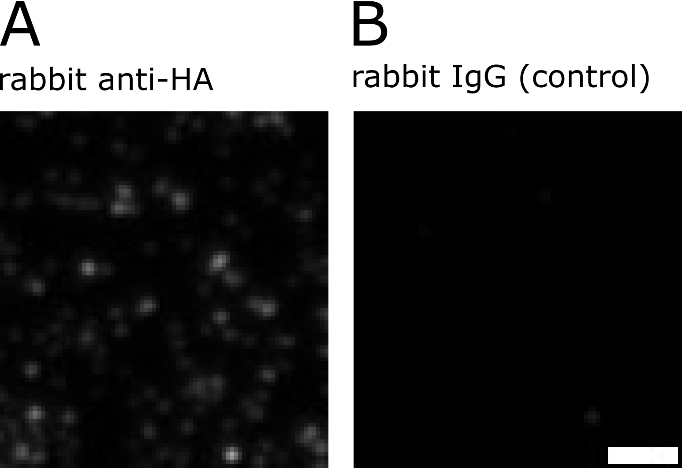

**Figure S3: Virus particles stained with anti-HA immunofluorescence and negative control.** A) Immobilized influenza particles were incubated with rabbit anti-HA IgG as primary ABs, followed by goat anti-rabbit IgG as secondary ABs. (see Fig 3) B) In the negative control, the primary ABs were exchanged for non-immunized rabbit IgGs. Scale Bar: 2 μm

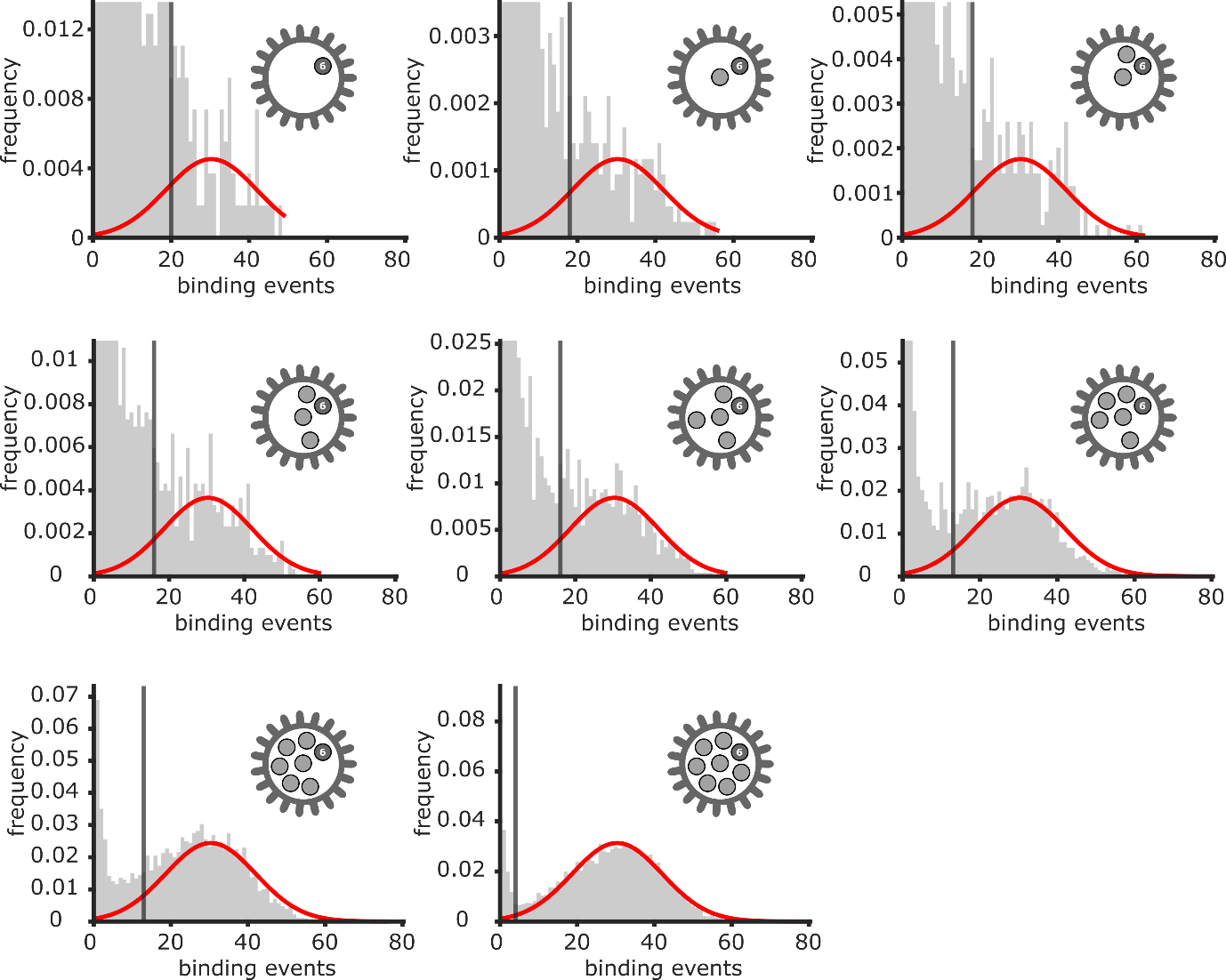

**Figure S4. The detection threshold of a segment (binding events) depends on the number of potential segments**, i.e. the presence of fluorescent traces in other detection rounds. The binding event population of NA (Fig S2) was divided according to how many fluorescent traces were found in imaging rounds of other segments, resulting in three populations, that were fitted with the Gaussian corresponding to the vRNP population (Fig S1, red curve), followed by determination of the treshold (black line). Due to the decreasing proportion of the background population compared to vRNP population, the detection threshold for particle candidates with more traces moves to lower values.

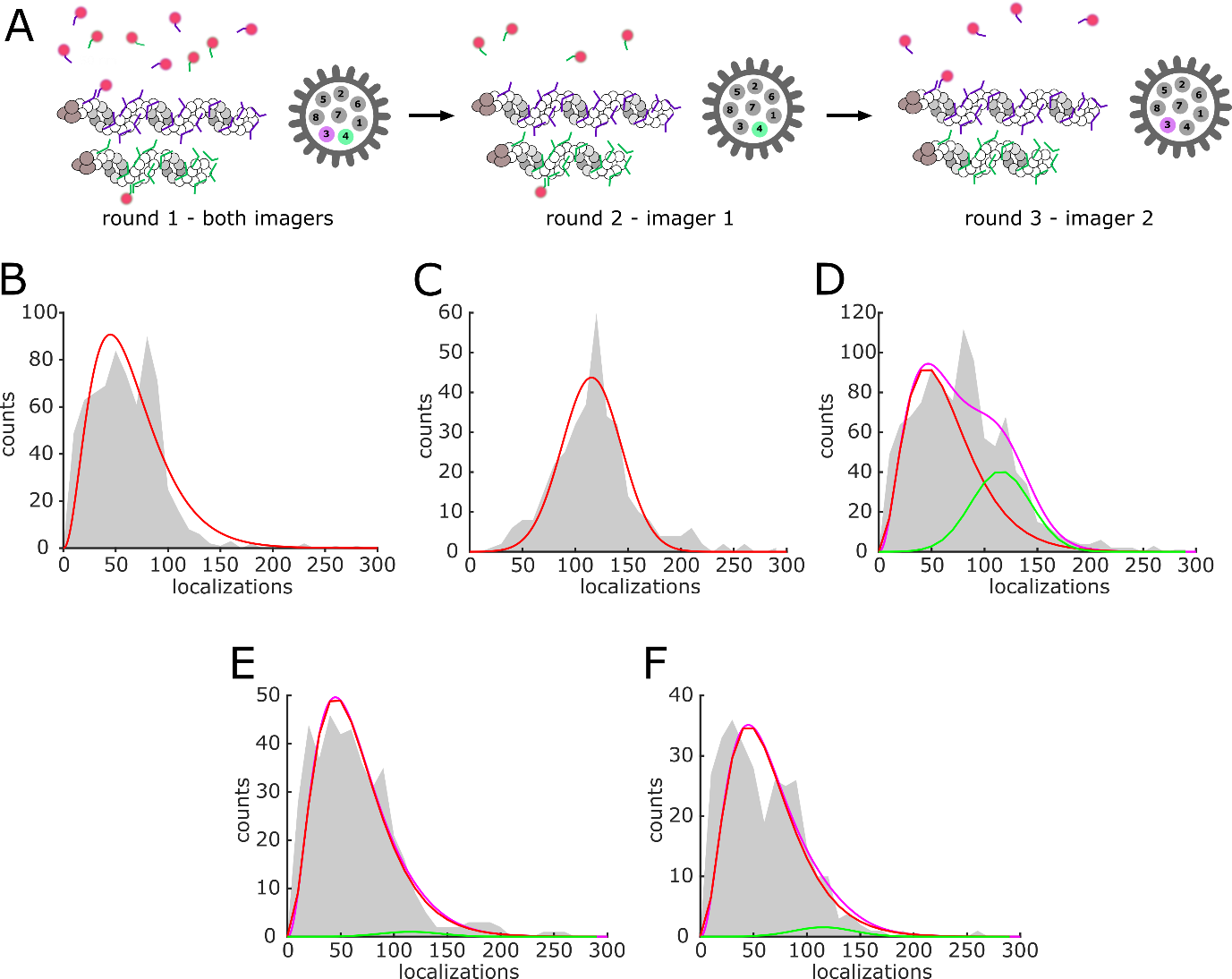

**Fig S5: Localization clusters in DNA-PAINT do generally not consist of more than one vRNP.** A) A typical DNA-PAINT experiment (Fig 1A) was conducted, where segments PA and HA were detected simultaneously by adding both assigned imager strands at once at a concentration, where binding events appeared separated (c = 5 nM, round 1). Subsequently both segments were detected individually to find out, whether the particles detected in round 1 posessed only HA, only PA or both vRNPs (rounds 2 and 3). Thus, we characterized the binding event distribution of one (B) and two segments (C) by fitting the one-segment population with a Gamma distribution and the two-segment population with a Normal distribution (red lines). D) When we fitted the entire population of PA/HA binding events (round one) with a mixed model comprising both distributions (mixed model: magenta, single vRNPs: red, double vRNPs: green) , we obtained a notable contribution from double segments - that is, particles containing both PA and HA) - as intended by th experimental design. E,F) When we fitted the binding events from round 2 and round 3 with the mixed model, the estimated contribution from double segments was less than 3.7 % (PA) and 8 % (HA), demonstrating that localization clusters on the surface predominantly depict individual vRNPs. The number of binding events in round 2 and 3 was slightly corrected for binding site loss by adjusting the number of localizations to fit the median localization number of the one-segment population in round 1.

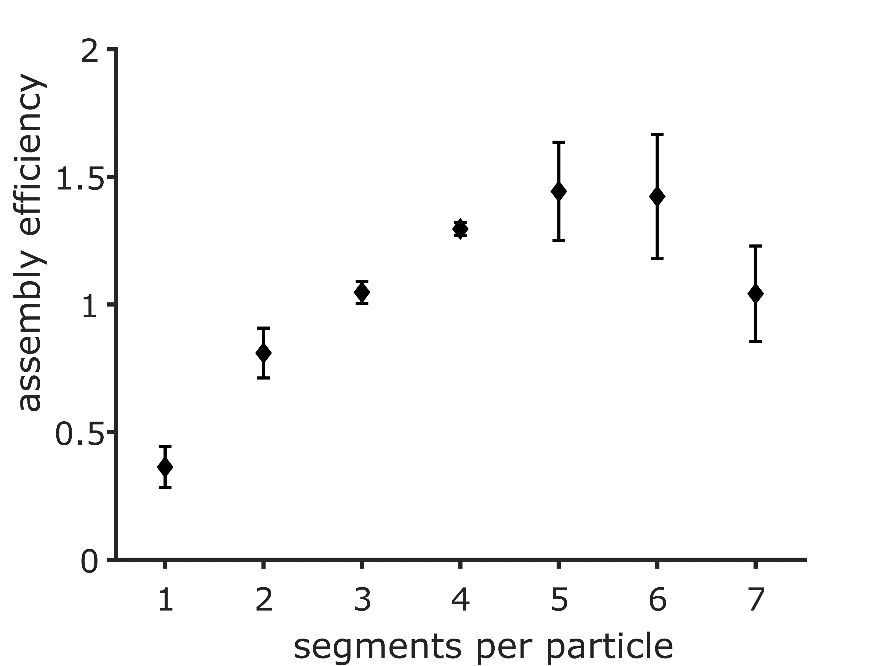

**Fig S6.** **Assembly efficiency (Eq. 2) for packaging intermediates (particles)** with a given number of segments. Error bars: STD of three technical replicates.

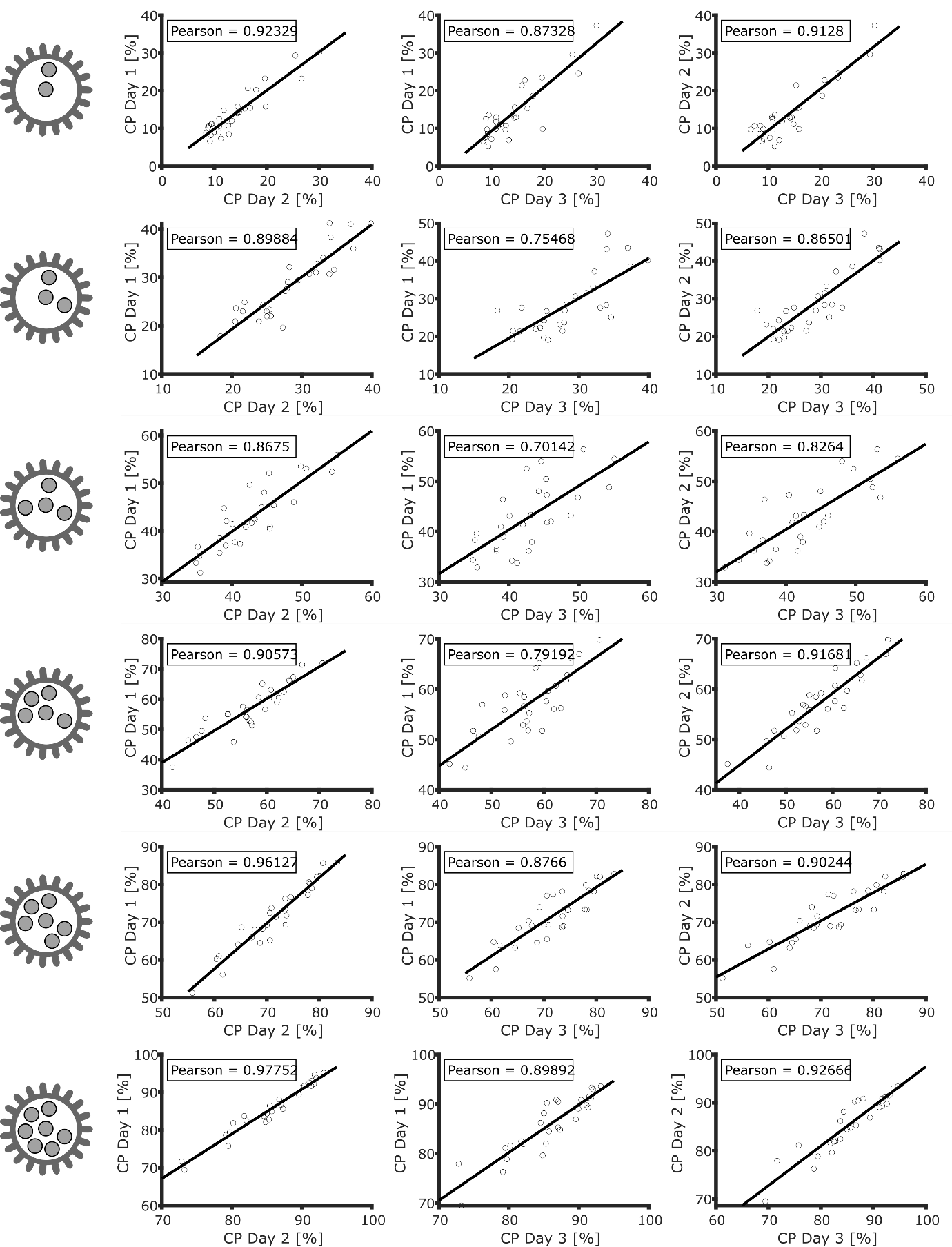

**Figure S7: Co-presence (CP) correlation between technical replicates (days) for particles with different segment counts** (left panel). Each of the data point in the plots represents one segment pair depicted in Fig 2C and Fig 2D (N = 28). The average Pearson coefficients for all segment counts ranged from 0.80 to 0.93.

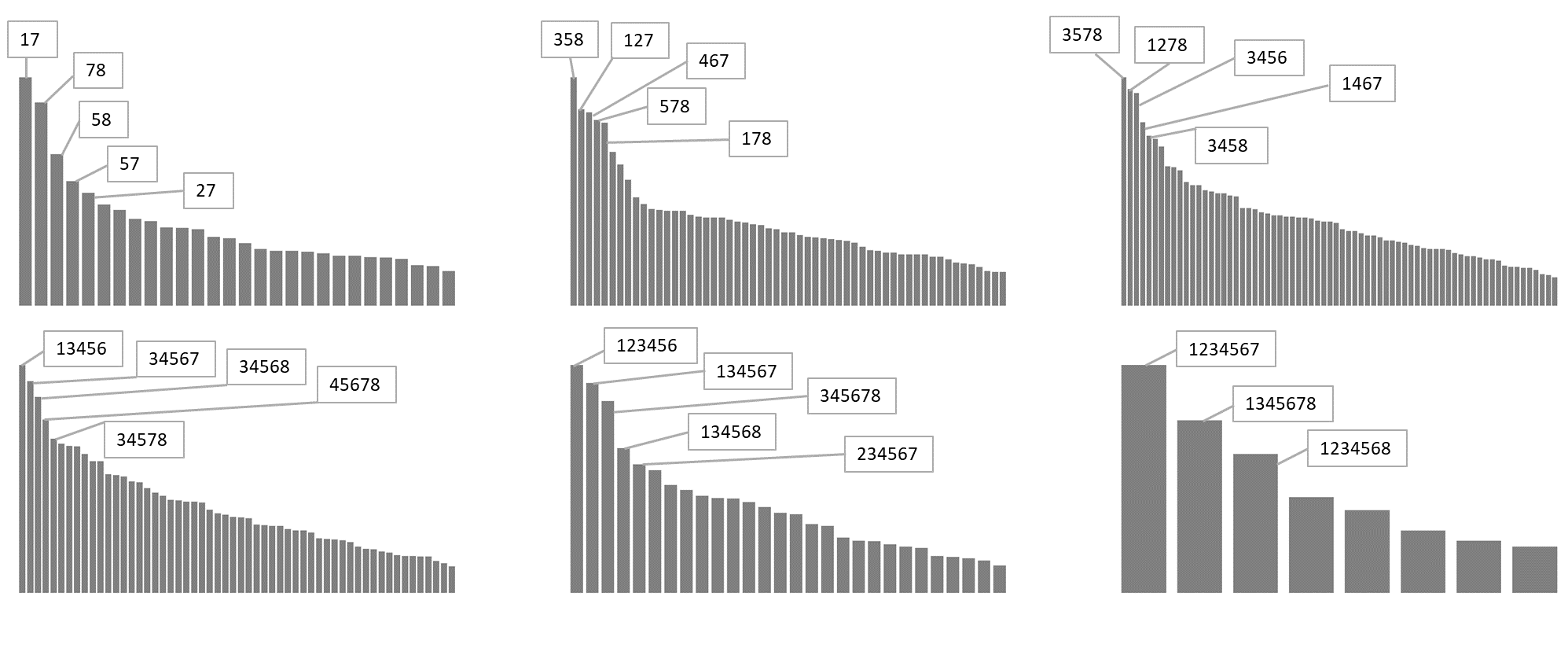

**Figure S8: Specific segment combinations for PR8 virus particles containing 2-7 segments.** Each subplot shows the total relative counts of segment combinations from particles with a given number of segments. See Haralampiev et al., Fig 4 for comparison with H3N2 (A/Panama) [47].

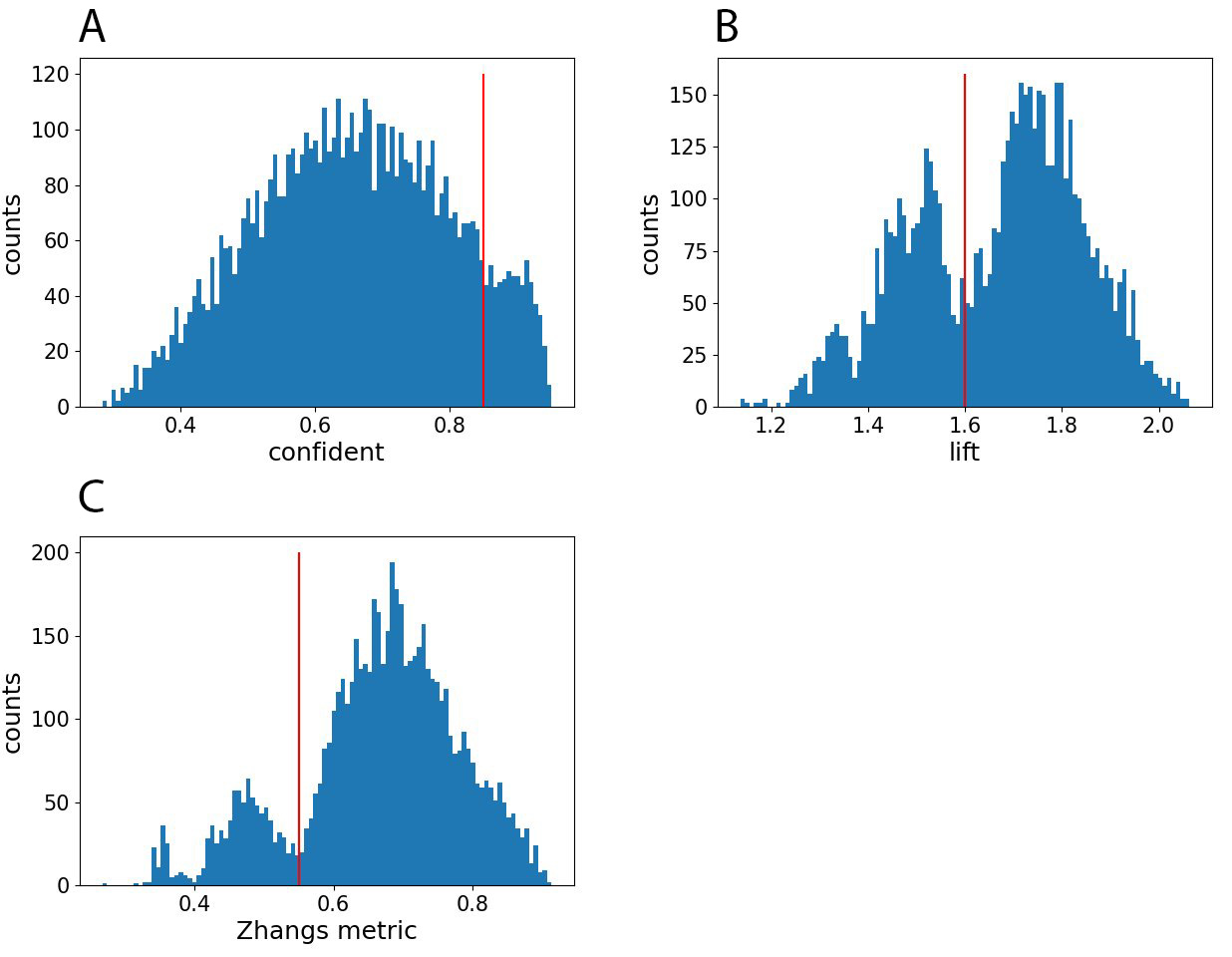

**Figure S9: Histogram of parameters used to filter association rules.** The red lines in the histogram indicate the thresholds applied for filtering. A) the histogram shows the confidence metric with a threshold of 0.85. B) the metric is lift with a threshold of 1.6. C) the histogram represents Zhang’s metric, with a threshold set at 0.55.

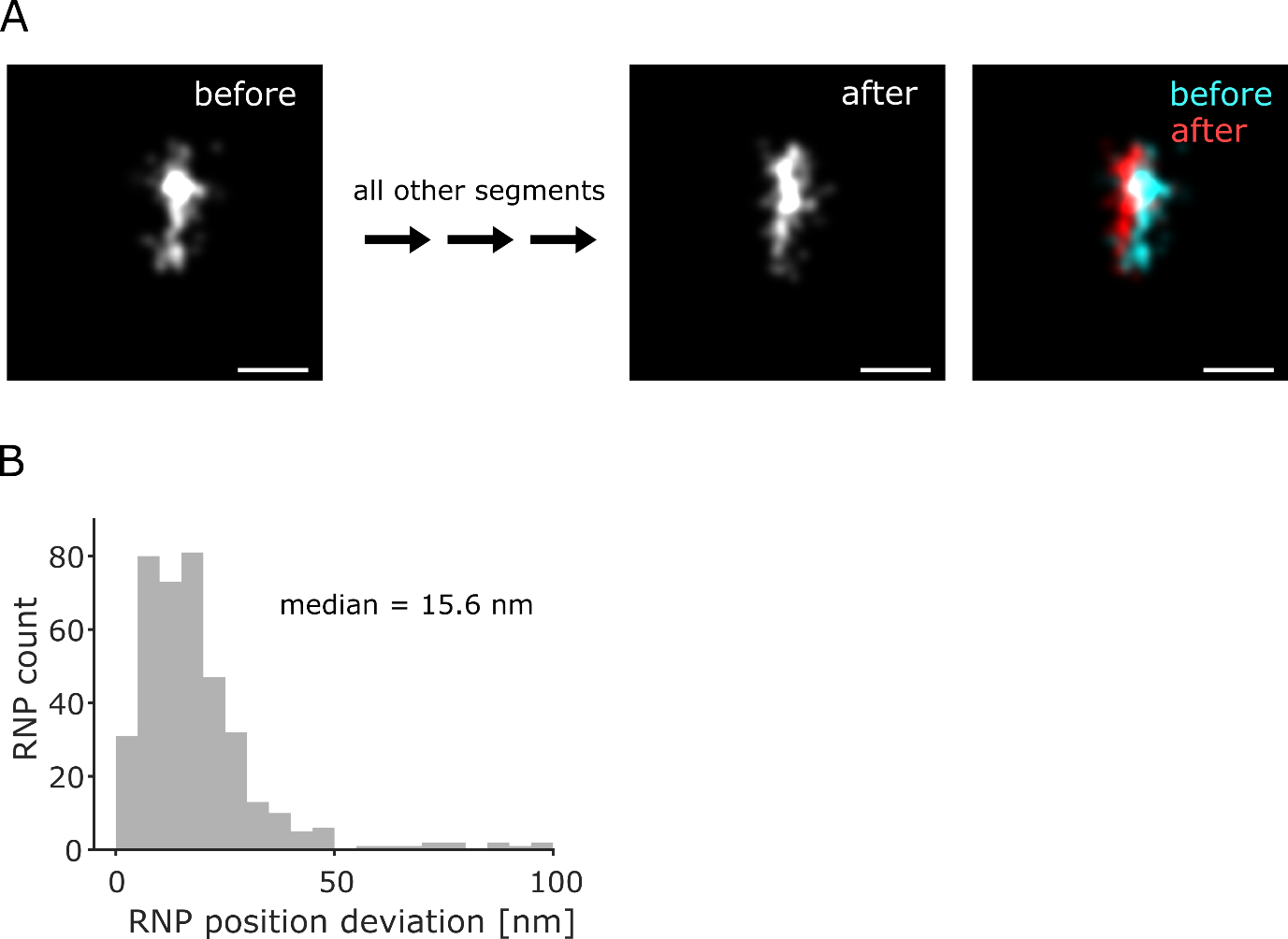

**Figure S10. Positional change of NA segments over the course of the imaging cycle** (8 imaging rounds, 24 liquid exchanges). A) Example of a super-resolved NA vRNP at the beginning (“before”) and the end of the imaging cycle (“after”). The composite on the right illustrates the positional shift. Scale bar: 50 nm. B) The mean shift in location of all NA vRNPs in a DNA-PAINT sample that could be detected both before and after imaging.

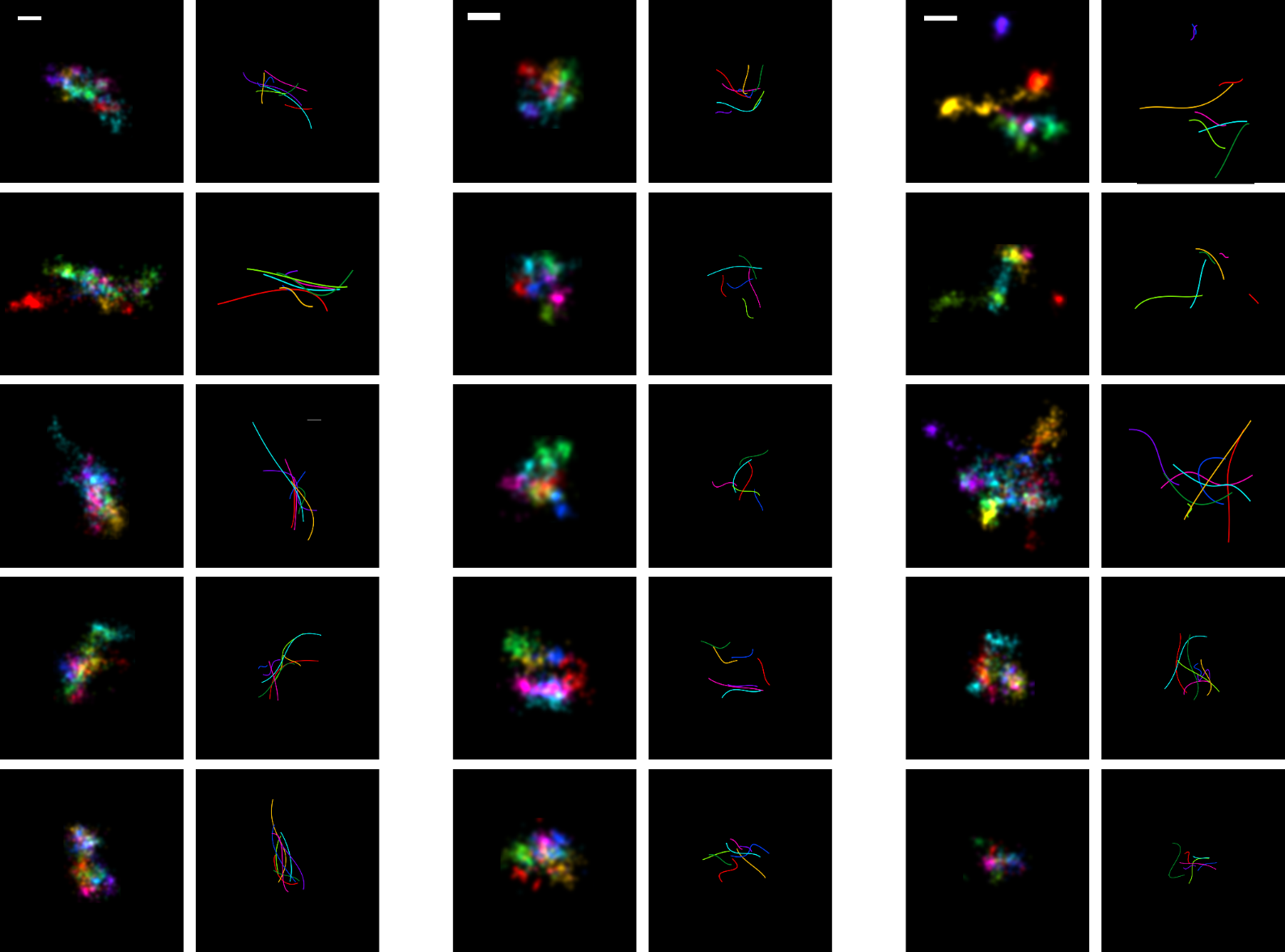

**Figure S11: Further examples of particle shapes as shown by the examples in Fig 4A**, including segment outlines. A) Elongated parallel vRNPs. B) Short, separated vRNPs. C) vRNPs with no apparent cardinal orientation or organization (disordered). Scale bars: 50 nm

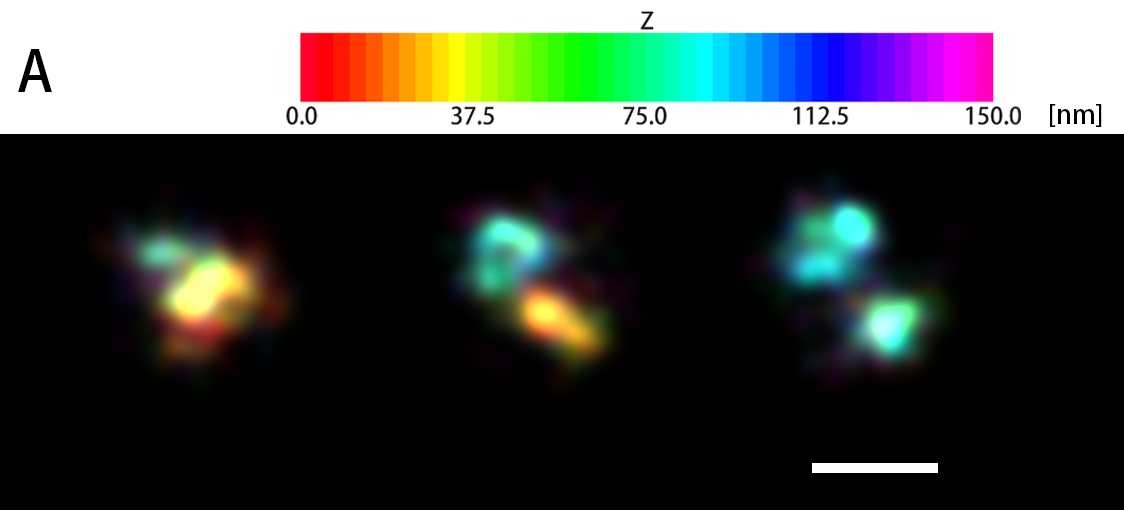

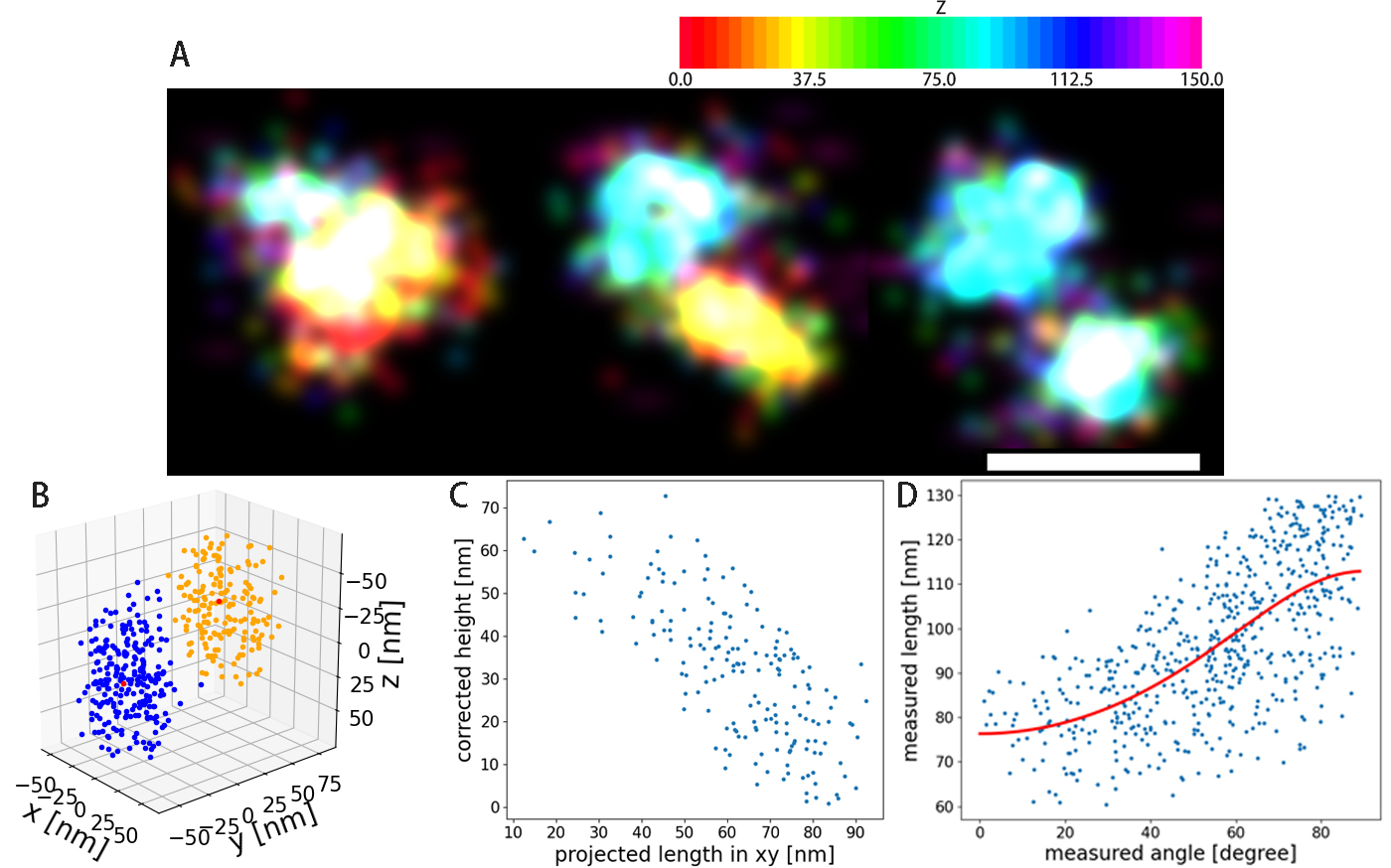

**Figure S12. Calibration and testing of the axial resolution using DNA origami.** A) DNA-PAINT images of 3D nano-rulers (GattaQuant) with variable orientations with respect to the xy plane. The z position is represented by the color of the localizations. The angle between nano-ruler and surface is decreasing from the left to the right panel. The scale bar is 100 nm. B) 3D plot of the localizations of the nano-ruler depicted in the middle panel of (A). The red dots represent the median positions of the localization clusters corresponding to the top (orange) and base (blue) of the nanoruler. C) Relationship between the height and length of the nanorulers, demonstrating different orientations with respect to the surface. D) Distribution of measured length and measured angle of the nano-rulers. Red line: model Fit of equation 1 (Materials and Methods) to determine the axial scaling factor.

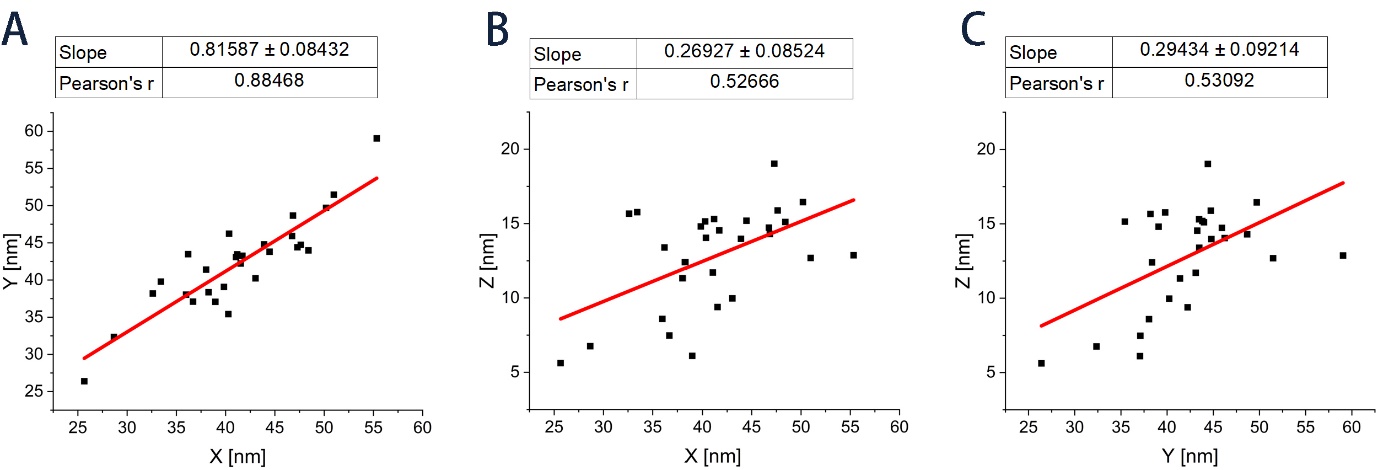

**Figure S13. The median of pairwise inter-segment distance along different directions.** (A) X and Y, (B) X and Z, (C) Y and Z. The red lines are the results of linear fits with intercept of zero. These distances are at least moderately correlated (PC 0.53 – 0.88). The values of the slopes in X-Y plane are significantly larger, showing that the median distances of segments in the xy -plane are longer (by a factor of ~ 3), compared to the median distances z direction.

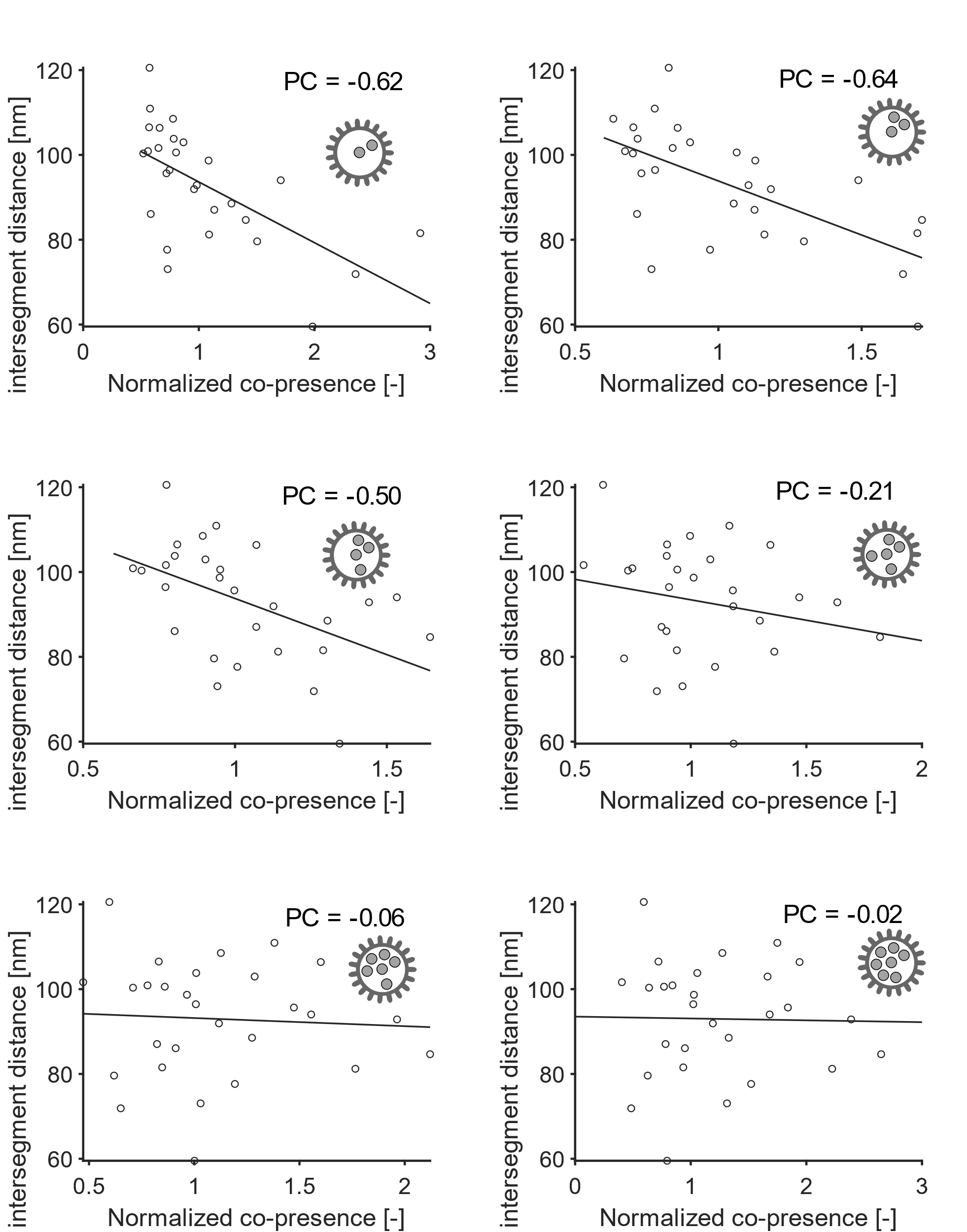

**Figure S14. Correlation between intersegment distances and normalized co-presence in particles with 2 – 7 segments** (compare Fig 5F). PC: Pearson’s coefficient.

**Table S1. Imaging conditions for each pair of imager and segment.** The imaging sequences are adapted from Schnitzbauer et al., 2017 [49]. Differences in sequence design are underlined.

| Imager | Target segment | Docking sequence | Imager sequence | Acquisition length (frames) | |
| --- | --- | --- | --- | --- | --- |
|  |  |  |  | Super-Res. | Segment-Stoch. |
| P1 | PB1 | TTATACATCTA | CTAGATGTAT-Dye | 4000 | 1000 |
| P2 | PB2 | TTATCTACATA | TATGTAGATC-Dye | 6700 | 1000 |
| P3 | NA | TTTCTTCATTA | GTAATGAAGA-Dye | 7000 | 750 |
| P4 | PA | TTATGAATCTA | GTAGATTCAT-Dye | 12300 | 900 |
| P5 | HA | TTTCAATGTAT | CATACATTGA-Dye | 12300 | 1000 |
| P6 | M | TTTTAGGTAAA | CTTTACCTAA-Dye | 17400 | 2250 |
| **P6L** | M | TTTTAGGTAAA | CTTTACCTAA**A**-Dye | 17400 | 2250 |
| P7 | NS | TTAATTGAGTA | GTACTCAATT-Dye | 12300 | 1300 |
| P8 | NP | TTATGTTAATG | CCATTAACAT-Dye | 8000 | 2250 |
| P9 | none | TTAATTAGGAT | CATCCTAATT-Dye |  |  |

**Table S2. Barcoded strands used in this study.**

| **Segment** | **Probe number** | **Hybridization Part (5' --> 3')** | **Imager Target (5' --> 3')** |
| --- | --- | --- | --- |
| PB2 | 1 | TTCGGATGGCCATCAATTAG | TTATCTACATA |
| PB2 | 2 | CATACTTACTGACAGCCAGA | TTATCTACATA |
| PB2 | 3 | AGACGTGGTGTTGGTAATGA | TTATCTACATA |
| PB2 | 4 | GGAGAGAAGGCTAATGTGCT | TTATCTACATA |
| PB2 | 5 | ATGAACTGAGCAACCTTGCG | TTATCTACATA |
| PB2 | 6 | AACAAGAGATATGGGCCAGC | TTATCTACATA |
| PB2 | 7 | ATTCCTCATTCTGGGCAAAG | TTATCTACATA |
| PB2 | 8 | TGTGAGGGGATCAGGAATGA | TTATCTACATA |
| PB2 | 9 | AGTTCTCCTCATTTACTGTG | TTATCTACATA |
| PB2 | 10 | GCTCCACCAAAGCAAAGTAG | TTATCTACATA |
| PB2 | 11 | ATAAAACTTCTTCCCTTCGC | TTATCTACATA |
| PB2 | 12 | GACATTTGATACCGCACAGA | TTATCTACATA |
| PB2 | 13 | TTAGAGGCCAATACAGTGGG | TTATCTACATA |
| PB2 | 14 | CAGTCTTTAGTACCTAAGGC | TTATCTACATA |
| PB2 | 15 | TCAGTGTTGGTCAATACCTA | TTATCTACATA |
| PB2 | 16 | CTCATCGTCAATGATGTGGG | TTATCTACATA |
| PB2 | 17 | AGGTCAGTGAAACACAGGGA | TTATCTACATA |
| PB2 | 18 | GAAATGTACTACTGTCTCCC | TTATCTACATA |
| PB2 | 19 | GACCGTTTTTTGAGAATCCG | TTATCTACATA |
| PB2 | 20 | TAGATGAGTACTCCAGCACG | TTATCTACATA |
| PB2 | 21 | TGACTCCAAGCATCGAGATG | TTATCTACATA |
| PB2 | 22 | GAATGATTGGGATATTGCCC | TTATCTACATA |
| PB2 | 23 | ATTGGGGAGTTGAACCTATC | TTATCTACATA |
| PB2 | 24 | ATAGGGCGAATCAGCGATTG | TTATCTACATA |
| PB2 | 25 | AAGCAGTCAGAGGTGATCTG | TTATCTACATA |
| PB2 | 26 | GCCATGGTATTTTCACAAGA | TTATCTACATA |
| PB2 | 27 | AGTCGATTGCCGAAGCAATA | TTATCTACATA |
| PB2 | 28 | GCTGATAGTGAGTGGGAGAG | TTATCTACATA |
| PB2 | 29 | GAGTTCACAATGGTTGGGAG | TTATCTACATA |
| PB2 | 30 | CTTCAGTTTTGGTGGATTCA | TTATCTACATA |
| PB2 | 31 | GCTGCAATGGGACTGAGAAT | TTATCTACATA |
| PB2 | 32 | CAACAGAAGAGCAAGCCGTG | TTATCTACATA |
| PB2 | 33 | GTAGACATCCTTAGGCAGAA | TTATCTACATA |
| PB2 | 34 | TTTATTGGAGATGTGCCACA | TTATCTACATA |
| PB2 | 35 | AGTATCAGCAGATCCACTAG | TTATCTACATA |
| PB2 | 36 | AGCTTGATTATTGCTGCTAG | TTATCTACATA |
| PB2 | 37 | ACTCAAGGAACATGCTGGGA | TTATCTACATA |
| PB2 | 38 | AGAGAGAACTGGTCCGCAAA | TTATCTACATA |
| PB2 | 39 | CCTTTGATGGTTGCATACAT | TTATCTACATA |
| PB2 | 40 | GAGCCAGGATACTAACATCG | TTATCTACATA |
| PB2 | 41 | GTTGTTTTCCCTAACGAAGT | TTATCTACATA |
| PB2 | 42 | GGCACAGGATGTAATCATGG | TTATCTACATA |
| PB2 | 43 | AAATCCTGGTCATGCAGATC | TTATCTACATA |
| PB2 | 44 | CCTGTCCATTTTAGAAACCA | TTATCTACATA |
| PB2 | 45 | AGGCTAAAGCATGGAACCTT | TTATCTACATA |
| PB2 | 46 | GCTGTGACATGGTGGAATAG | TTATCTACATA |
| PB2 | 47 | ATCAGACCGAGTGATGGTAT | TTATCTACATA |
| PB2 | 48 | CTCACAAAAACCACCGTGGA | TTATCTACATA |
| PB1 | 1 | AGTGAATTTAGCTTGTCCTT | TTATACATCTA |
| PB1 | 2 | TTGAAGAGCTCAGACGGCAA | TTATACATCTA |
| PB1 | 3 | CCGAATTGATGCACGGATTG | TTATACATCTA |
| PB1 | 4 | CAGAAGACCAGTCGGGATAT | TTATACATCTA |
| PB1 | 5 | GGAGTATGATGCTGTTGCAA | TTATACATCTA |
| PB1 | 6 | CACATGGTCCAGCCAAAAAC | TTATACATCTA |
| PB1 | 7 | GTTTATGCAACCCACTGAAC | TTATACATCTA |
| PB1 | 8 | ATGGATGAGGATTACCAGGG | TTATACATCTA |
| PB1 | 9 | ATTCCTGAAGTCTGCCTAAA | TTATACATCTA |
| PB1 | 10 | ACGGAGGCCCAAATTTATAC | TTATACATCTA |
| PB1 | 11 | TACAGGTACACGTACCGATG | TTATACATCTA |
| PB1 | 12 | CCCTTCAGTTGTTCATCAAA | TTATACATCTA |
| PB1 | 13 | AATGATCTTGGTCCAGCAAC | TTATACATCTA |
| PB1 | 14 | AACGAGTCAGCGGACATGAG | TTATACATCTA |
| PB1 | 15 | TTGTTGCCAATTTCAGCATG | TTATACATCTA |
| PB1 | 16 | CGGAGTCGACAGGTTTTATC | TTATACATCTA |
| PB1 | 17 | CACCCAATCATGAAGGGATT | TTATACATCTA |
| PB1 | 18 | CAATCCTCTGACGATTTTGC | TTATACATCTA |
| PB1 | 19 | CCAAGACTACTTACTGGTGG | TTATACATCTA |
| PB1 | 20 | TAAGCACTGTATTAGGCGTC | TTATACATCTA |
| PB1 | 21 | GGAATGATGATGGGCATGTT | TTATACATCTA |
| PB1 | 22 | TCGCTCTTAATAGAGGGGAC | TTATACATCTA |
| PB1 | 23 | ACCTGCAGAAATGCTAGCAA | TTATACATCTA |
| PB1 | 24 | GCGAGACTGGGAAAAGGGTA | TTATACATCTA |
| PB1 | 25 | ATGTTCTAAGTATTGCTCCA | TTATACATCTA |
| PB1 | 26 | ATATGACCAGAAATCAGCCC | TTATACATCTA |
| PB1 | 27 | GAATCCTCGGATGTTTTTGG | TTATACATCTA |
| PB1 | 28 | ACTTTCTTTCACCATCACTG | TTATACATCTA |
| PB1 | 29 | GATGACCAATTCTCAGGACA | TTATACATCTA |
| PB1 | 30 | TGAACAATCAGGGTTGCCAG | TTATACATCTA |
| PB1 | 31 | TGAGACACTGGCAAGGAGTA | TTATACATCTA |
| PB1 | 32 | AGGGATGCAAATAAGGGGGT | TTATACATCTA |
| PB1 | 33 | AACGGAGAGCAATTGCAACC | TTATACATCTA |
| PB1 | 34 | AGATGCTGAGAGAGGGAAGC | TTATACATCTA |
| PB1 | 35 | TTAGAGCATTGACCCTGAAC | TTATACATCTA |
| PB1 | 36 | GGCTCATAGACTTCCTTAAG | TTATACATCTA |
| PB1 | 37 | TGTTCAGATCAAATGGCCTC | TTATACATCTA |
| PB1 | 38 | GCATTGGCCAACACAATAGA | TTATACATCTA |
| PB1 | 39 | TAGAAACCAACCTGCTGCAA | TTATACATCTA |
| PB1 | 40 | AGACCTATGACTGGACTCTA | TTATACATCTA |
| PB1 | 41 | GGAGGTTGTTCAGCAAACAC | TTATACATCTA |
| PB1 | 42 | GATTGTGTATTGGAGGCGAT | TTATACATCTA |
| PB1 | 43 | ATGAACCAAGTGGTTATGCC | TTATACATCTA |
| PB1 | 44 | CAACTCAACCCGATTGATGG | TTATACATCTA |
| PB1 | 45 | AGCACAACTTTCCCTTATAC | TTATACATCTA |
| PB1 | 46 | AAGTGCCAGCACAAAATGCT | TTATACATCTA |
| PB1 | 47 | TCAATCCGACCTTACTTTTC | TTATACATCTA |
| PB1 | 48 | GCGAAAGCAGGCAAACCATT | TTATACATCTA |
| PA | 1 | TGCTATCCATACTGTCCAAA | TTATGAATCTA |
| PA | 2 | ATTGAGTTAGTTGTGGCAGT | TTATGAATCTA |
| PA | 3 | GGTTCAACTCATTCCTTACA | TTATGAATCTA |
| PA | 4 | TGGGTTTTGCTTAATGCTTC | TTATGAATCTA |
| PA | 5 | GAGGAGTGCCTGATTAATGA | TTATGAATCTA |
| PA | 6 | ATCTTGGGGGGCTATATGAA | TTATGAATCTA |
| PA | 7 | CTTATCGTTCAGGCTCTTAG | TTATGAATCTA |
| PA | 8 | GCATCTCCACAACTAGAAGG | TTATGAATCTA |
| PA | 9 | AGTCGGTATTCAACAGCTTG | TTATGAATCTA |
| PA | 10 | GGTCTGCAGGACTTTATTAG | TTATGAATCTA |
| PA | 11 | GTGGAGGAAAGTTCCATTGG | TTATGAATCTA |
| PA | 12 | ATCAGAAACATGGCCCATTG | TTATGAATCTA |
| PA | 13 | CTGAGTCCTCTGTCAAAGAG | TTATGAATCTA |
| PA | 14 | CCTCCAGTCACTTCAACAAA | TTATGAATCTA |
| PA | 15 | CCATGTTCTTGTATGTGAGA | TTATGAATCTA |
| PA | 16 | TATAAGAAGTGCCATAGGCC | TTATGAATCTA |
| PA | 17 | TGACCCAAGACTTGAACCAC | TTATGAATCTA |
| PA | 18 | AGGAAGATCCCACTTAAGGA | TTATGAATCTA |
| PA | 19 | GACCAACTTGTATGGTTTCA | TTATGAATCTA |
| PA | 20 | GAACTAAGGAGGGAAGGCGA | TTATGAATCTA |
| PA | 21 | TTGCTTAATGCATCTTGTGC | TTATGAATCTA |
| PA | 22 | GGGGGTGTACATCAATACTG | TTATGAATCTA |
| PA | 23 | TTTCACATCAGAGGTGTCTC | TTATGAATCTA |
| PA | 24 | TTCAAGCTGGATAGAGCTCG | TTATGAATCTA |
| PA | 25 | ACAAGGCATGCGAACTGACA | TTATGAATCTA |
| PA | 26 | GGCACCAGAAAAGGTAGACT | TTATGAATCTA |
| PA | 27 | AAGTGGGCACTTGGTGAGAA | TTATGAATCTA |
| PA | 28 | CTTCTGTCATGGAAGCAAGT | TTATGAATCTA |
| PA | 29 | ACCGCTATATGATGCAATCA | TTATGAATCTA |
| PA | 30 | GTCCAAATTCCTACTGATGG | TTATGAATCTA |
| PA | 31 | CACTTAGACTTCCGAATGGG | TTATGAATCTA |
| PA | 32 | CAAGCTGTCTCAAATGTCCA | TTATGAATCTA |
| PA | 33 | GAACCGAACGGCTACATTGA | TTATGAATCTA |
| PA | 34 | TAGAGCCTATGTGGATGGAT | TTATGAATCTA |
| PA | 35 | ACTTCTCCAGCCTTGAAAAT | TTATGAATCTA |
| PA | 36 | CACAGGAACAATGCGCAAGC | TTATGAATCTA |
| PA | 37 | GGGCTAGGATCAAAACCAGA | TTATGAATCTA |
| PA | 38 | CCACAAAGGCAGACTACACT | TTATGAATCTA |
| PA | 39 | TCTCGTTCACTGGGGAAGAA | TTATGAATCTA |
| PA | 40 | GGGGCTGAGAAACCAAAGTT | TTATGAATCTA |
| PA | 41 | TGGCCTGGACAGTAGTAAAC | TTATGAATCTA |
| PA | 42 | ATCGAGGGAAGAGATCGCAC | TTATGAATCTA |
| PA | 43 | GCAAGGCGAGTCAATAATCG | TTATGAATCTA |
| PA | 44 | TTGCAGCAATATGCACTCAC | TTATGAATCTA |
| PA | 45 | GAAAGAGTATGGGGAGGACC | TTATGAATCTA |
| PA | 46 | GATTGTCGAGCTTGCGGAAA | TTATGAATCTA |
| PA | 47 | TTTTGTGCGACAATGCTTCA | TTATGAATCTA |
| PA | 48 | CGAAAGCAGGTACTGATCCA | TTATGAATCTA |
| HA | 1 | TGCAGAATATGCATCTGAGA | TTTCAATGTAT |
| HA | 2 | GTGTTCTAATGGATCTTTGC | TTTCAATGTAT |
| HA | 3 | CTGGGGGCAATCAGTTTCTG | TTTCAATGTAT |
| HA | 4 | CAGTTCACTGGTGCTTTTGG | TTTCAATGTAT |
| HA | 5 | TGGCGATCTACTCAACTGTC | TTTCAATGTAT |
| HA | 6 | ATCAATGGGGATCTATCAGA | TTTCAATGTAT |
| HA | 7 | GTGTGACAATGAATGCATGG | TTTCAATGTAT |
| HA | 8 | GGATGTTTTGAGTTCTACCA | TTTCAATGTAT |
| HA | 9 | TCTGGATTTCCATGACTCAA | TTTCAATGTAT |
| HA | 10 | GGATTTCTGGACATTTGGAC | TTTCAATGTAT |
| HA | 11 | GGTGAACACTGTTATCGAGA | TTTCAATGTAT |
| HA | 12 | TGCCATTAACGGGATTACAA | TTTCAATGTAT |
| HA | 13 | TTGCCGGTTTTATTGAAGGG | TTTCAATGTAT |
| HA | 14 | TCCAGAGGTCTATTTGGAGC | TTTCAATGTAT |
| HA | 15 | AAGGAACAATCCGTCCATTC | TTTCAATGTAT |
| HA | 16 | AATTGAGGATGGTTACAGGA | TTTCAATGTAT |
| HA | 17 | CCCAAAATACGTCAGGAGTG | TTTCAATGTAT |
| HA | 18 | ACCCAGTCACAATAGGAGAG | TTTCAATGTAT |
| HA | 19 | AGTCTCCCTTACCAGAATAT | TTTCAATGTAT |
| HA | 20 | ACCCCTGGGAGCTATAAACA | TTTCAATGTAT |
| HA | 21 | GCATGAGTGTAACACGAAGT | TTTCAATGTAT |
| HA | 22 | ATCATCACCTCAAACGCATC | TTTCAATGTAT |
| HA | 23 | CACTGAGTAGAGGCTTTGGG | TTTCAATGTAT |
| HA | 24 | ATAGCACCAATGTATGCTTT | TTTCAATGTAT |
| HA | 25 | ACTATTACTGGACCTTGCTA | TTTCAATGTAT |
| HA | 26 | AGAGATCAAGCTGGGAGGAT | TTTCAATGTAT |
| HA | 27 | CCCCGGAAATAGCAGAAAGA | TTTCAATGTAT |
| HA | 28 | GAAAATGCTTATGTCTCTGT | TTTCAATGTAT |
| HA | 29 | CGCCTAACAGTAAGGAACAA | TTTCAATGTAT |
| HA | 30 | CGGAGAAGGAGGGCTCATAC | TTTCAATGTAT |
| HA | 31 | TACAGAAATTTGCTATGGCT | TTTCAATGTAT |
| HA | 32 | CCATGAGGGGAAAAGCAGTT | TTTCAATGTAT |
| HA | 33 | ACAAACGGAGTAACGGCAGC | TTTCAATGTAT |
| HA | 34 | AAAGAAAGCTCATGGCCCAA | TTTCAATGTAT |
| HA | 35 | ATTGAGCTCAGTGTCATCAT | TTTCAATGTAT |
| HA | 36 | CATCGACTATGAGGAGCTGA | TTTCAATGTAT |
| HA | 37 | CCTACATTGTAGAAACACCA | TTTCAATGTAT |
| HA | 38 | CTGCTTCCAGTGAGATCATG | TTTCAATGTAT |
| HA | 39 | CTCTTGGGAAACCCAGAATG | TTTCAATGTAT |
| HA | 40 | GGAAATGTAACATCGCCGGA | TTTCAATGTAT |
| HA | 41 | AAGGAATAGCCCCACTACAA | TTTCAATGTAT |
| HA | 42 | GCCACAACGGAAAACTATGT | TTTCAATGTAT |
| HA | 43 | TCTGTTAACCTGCTCGAAGA | TTTCAATGTAT |
| HA | 44 | CTCGAGAAGAATGTGACAGT | TTTCAATGTAT |
| HA | 45 | TCAACCGACACTGTTGACAC | TTTCAATGTAT |
| HA | 46 | TATAGGCTACCATGCGAACA | TTTCAATGTAT |
| HA | 47 | GCTGCAGATGCAGACACAAT | TTTCAATGTAT |
| HA | 48 | TGAAGGCAAACCTACTGGTC | TTTCAATGTAT |
| NP | 1 | CGAGCTCTCGGACGAAAAGG | TTATGTTAATG |
| NP | 2 | AGAAGATGTGTCTTTCCAGG | TTATGTTAATG |
| NP | 3 | CATCTGACATGAGGACCGAA | TTATGTTAATG |
| NP | 4 | TGGCAGCATTCAATGGGAAT | TTATGTTAATG |
| NP | 5 | ATCTCCCTTTTGACAGAACA | TTATGTTAATG |
| NP | 6 | TCAGCATACAACCTACGTTC | TTATGTTAATG |
| NP | 7 | AATCAACAGAGGGCATCTGC | TTATGTTAATG |
| NP | 8 | CCATAAGGACCAGAAGTGGA | TTATGTTAATG |
| NP | 9 | AAGAGGGAAGCTTTCCACTA | TTATGTTAATG |
| NP | 10 | TCAAAGGGACGAAGGTGCTC | TTATGTTAATG |
| NP | 11 | ATTCTGCCGCATTTGAAGAT | TTATGTTAATG |
| NP | 12 | TCCAGCACACAAGAGTCAAC | TTATGTTAATG |
| NP | 13 | GTACAGCCTAATCAGACCAA | TTATGTTAATG |
| NP | 14 | CAGACTGCTTCAAAACAGCC | TTATGTTAATG |
| NP | 15 | CTCTCTAGTCGGAATAGACC | TTATGTTAATG |
| NP | 16 | CTGCACTCATATTGAGAGGG | TTATGTTAATG |
| NP | 17 | GATCTCACTTTTCTAGCACG | TTATGTTAATG |
| NP | 18 | GATCAAGTGAGAGAGAGCCG | TTATGTTAATG |
| NP | 19 | AACTTCTGGAGGGGTGAGAA | TTATGTTAATG |
| NP | 20 | CTCTCTGATGCAAGGTTCAA | TTATGTTAATG |
| NP | 21 | GAGGACAAGAGCTCTTGTTC | TTATGTTAATG |
| NP | 22 | CTCACATGATGATCTGGCAT | TTATGTTAATG |
| NP | 23 | CTGGCGCCAAGCTAATAATG | TTATGTTAATG |
| NP | 24 | CTGGAGGACCTATATACAGG | TTATGTTAATG |
| NP | 25 | TGCGGGGAAAGATCCTAAGA | TTATGTTAATG |
| NP | 26 | AGAGAGAATGGTGCTCTCTG | TTATGTTAATG |
| NP | 27 | GGGACGGTTGATCCAAAACA | TTATGTTAATG |
| NP | 28 | TGTGCACCGAACTCAAACTC | TTATGTTAATG |
| NP | 29 | ATTGGACGATTCTACATCCA | TTATGTTAATG |
| NP | 30 | TGAAATCAGAGCATCCGTCG | TTATGTTAATG |
| NP | 31 | ACTGATGGAGAACGCCAGAA | TTATGTTAATG |
| NP | 32 | TCTCAAGGCACCAAACGATC | TTATGTTAATG |
| NP | 33 | TCACTCACTGAGTGACATCA | TTATGTTAATG |
| NA | 1 | GTCTGTTCAAAAAACTCCTT | TTTCTTCATTA |
| NA | 2 | TGCCATTCAGCATTGACAAG | TTTCTTCATTA |
| NA | 3 | TACTGTAGATTGGTCTTGGC | TTTCTTCATTA |
| NA | 4 | CTAGACTGTATGAGGCCGTG | TTTCTTCATTA |
| NA | 5 | TCAACATCCTGAGCTGACAG | TTTCTTCATTA |
| NA | 6 | TCAGGGTATAGCGGAAGTTT | TTTCTTCATTA |
| NA | 7 | GATGTTGTGGCAATGACTGA | TTTCTTCATTA |
| NA | 8 | TAGTAAGTTCTCTGTGAGGC | TTTCTTCATTA |
| NA | 9 | CCTAATGGATGGACAGAGAC | TTTCTTCATTA |
| NA | 10 | TGGGTTTGAGATGATTTGGG | TTTCTTCATTA |
| NA | 11 | GACCAAAAGTCACAGTTCCA | TTTCTTCATTA |
| NA | 12 | GGTATGGTAATGGTGTTTGG | TTTCTTCATTA |
| NA | 13 | GGAGCAAACGGAGTAAAGGG | TTTCTTCATTA |
| NA | 14 | TAGGATACATCTGCAGTGGG | TTTCTTCATTA |
| NA | 15 | CATGGGTGTCTTTCGATCAA | TTTCTTCATTA |
| NA | 16 | AATTGGCATGGTTCGAACCG | TTTCTTCATTA |
| NA | 17 | TTACCCTGATACCGGCAAAG | TTTCTTCATTA |
| NA | 18 | CTCACTATGAGGAATGTTCC | TTTCTTCATTA |
| NA | 19 | CGATAGAGTTGAATGCACCT | TTTCTTCATTA |
| NA | 20 | TCGAAAAGGGGAAGGTTACT | TTTCTTCATTA |
| NA | 21 | GCTGGCCTCGTACAAAATTT | TTTCTTCATTA |
| NA | 22 | ACTATAATGACTGATGGCCC | TTTCTTCATTA |
| NA | 23 | GCCTGTGTAAATGGTTCATG | TTTCTTCATTA |
| NA | 24 | TTGAGGACACAAGAGTCTGA | TTTCTTCATTA |
| NA | 25 | TAATGGAGCAGTGGCTGTAT | TTTCTTCATTA |
| NA | 26 | CAATCGGAATTTCAGGTCCA | TTTCTTCATTA |
| NA | 27 | TTGGTCAGCAAGTGCATGTC | TTTCTTCATTA |
| NA | 28 | TATAGGGCCTTAATGAGCTG | TTTCTTCATTA |
| NA | 29 | GTGGGACTGTTAAGGACAGA | TTTCTTCATTA |
| NA | 30 | GCCTTACTGAATGACAAGCA | TTTCTTCATTA |
| NA | 31 | GAATGCAGGACCTTTTTTCT | TTTCTTCATTA |
| NA | 32 | TTTGTCATAAGAGAGCCCTT | TTTCTTCATTA |
| NA | 33 | AATTGGTTCCAAAGGAGACG | TTTCTTCATTA |
| NA | 34 | GGTGGGCTATATACAGCAAA | TTTCTTCATTA |
| NA | 35 | ATTCATCTCTTTGTCCCATC | TTTCTTCATTA |
| NA | 36 | ACTTCAGTGATATTAACCGG | TTTCTTCATTA |
| NA | 37 | ATAGCACCTGGGTAAAGGAC | TTTCTTCATTA |
| NA | 38 | GCAACCAAAACATCATTACC | TTTCTTCATTA |
| NA | 39 | GGAAGTCAAAACCATACTGG | TTTCTTCATTA |
| NA | 40 | TCTCAATATGGATTAGCCAT | TTTCTTCATTA |
| NA | 41 | AGTCGGACTAATTAGCCTAA | TTTCTTCATTA |
| NA | 42 | CCATTGGATCAATCTGTCTG | TTTCTTCATTA |
| M | 1 | GATGGTCATTTTGTCAGCAT | TTTTAGGTAAA |
| M | 2 | AAGGAACAGCAGAGTGCTGT | TTTTAGGTAAA |
| M | 3 | TACGGAAGGAGTGCCAAAGT | TTTTAGGTAAA |
| M | 4 | CCGTCGCTTTAAATACGGAC | TTTTAGGTAAA |
| M | 5 | GCACTTGACATTGTGGATTC | TTTTAGGTAAA |
| M | 6 | TTGCCGCAAATATCATTGGG | TTTTAGGTAAA |
| M | 7 | TCAGAAACGAATGGGGGTGC | TTTTAGGTAAA |
| M | 8 | TCCAGTGCTGGTCTGAAAAA | TTTTAGGTAAA |
| M | 9 | AACCATTGGAACTCATCCTA | TTTTAGGTAAA |
| M | 10 | TCAGGCTAGACAAATGGTGC | TTTTAGGTAAA |
| M | 11 | CAAATGGCTGGATCGAGTGA | TTTTAGGTAAA |
| M | 12 | CACTACAGCTAAGGCTATGG | TTTTAGGTAAA |
| M | 13 | GGTGACAACAACCAATCCAC | TTTTAGGTAAA |
| M | 14 | GTATGTGCAACCTGTGAACA | TTTTAGGTAAA |
| M | 15 | GTATGGGCCTCATATACAAC | TTTTAGGTAAA |
| M | 16 | ATCTCACTCAGTTATTCTGC | TTTTAGGTAAA |
| M | 17 | TTAGGATTTGTGTTCACGCT | TTTTAGGTAAA |
| M | 18 | AGACAAGACCAATCCTGTCA | TTTTAGGTAAA |
| M | 19 | GATCTTGAGGTTCTCATGGA | TTTTAGGTAAA |
| M | 20 | TGAAGATGTCTTTGCAGGGA | TTTTAGGTAAA |
| M | 21 | GGTCGAAACGTACGTACTCT | TTTTAGGTAAA |
| NS | 1 | GCTTGAAGTGGAGCAAGAGA | TTAATTGAGTA |
| NS | 2 | TTTATGCAAGCCTTACATCT | TTAATTGAGTA |
| NS | 3 | TCCACTCACTCCAAAACAGA | TTAATTGAGTA |
| NS | 4 | TCTACAGAGATTCGCTTGGA | TTAATTGAGTA |
| NS | 5 | AACACAGTTCGAGTCTCTGA | TTAATTGAGTA |
| NS | 6 | AAAATGCAGTTGGAGTCCTC | TTAATTGAGTA |
| NS | 7 | CAGGACATACTGCTGAGGAT | TTAATTGAGTA |
| NS | 8 | AAATTTCACCATTGCCTTCT | TTAATTGAGTA |
| NS | 9 | CACCGAAGAGGGAGCAATTG | TTAATTGAGTA |
| NS | 10 | CTCTAATATTGCTAAGGGCT | TTAATTGAGTA |
| NS | 11 | TCAGTGTGATTTTTGACCGG | TTAATTGAGTA |
| NS | 12 | ACCAGGCGATCATGGATAAG | TTAATTGAGTA |
| NS | 13 | AAATGTCAAGGGACTGGTCC | TTAATTGAGTA |
| NS | 14 | GCGTTACCTAACTGACATGA | TTAATTGAGTA |
| NS | 15 | GAAGAATCCGATGAGGCACT | TTAATTGAGTA |
| NS | 16 | CGTGCTGGAAAGCAGATAGT | TTAATTGAGTA |
| NS | 17 | CTTCGCCGAGATCAGAAATC | TTAATTGAGTA |
| NS | 18 | GCAGACCAAGAACTAGGTGA | TTAATTGAGTA |
| NS | 19 | TGGATCCAAACACTGTGTCA | TTAATTGAGTA |

**Table S3. Imaging conditions for super-resolution and high-throughput stoichiometry experiments in comparison.**

|  |  | super-resolution | segment stoichiometry |
| --- | --- | --- | --- |
| Imager concentration | | 5 nM | 50 nM |
| Exposure time | | 200 ms | 20 ms |
| Total imaging time | frames | 4000 -17400 | 750-2250 |
|  | minutes | 13-58 | 0.25-0.75 |
| Sample size (FOVs) | | 1 | 49 |
